# Structural determinants of persulfide-sensing specificity in a dithiol-based transcriptional regulator

**DOI:** 10.1101/2020.03.22.001966

**Authors:** Daiana A. Capdevila, Brenna J. C. Walsh, Yifan Zhang, Christopher Dietrich, Giovanni Gonzalez-Gutierrez, David P. Giedroc

## Abstract

Cysteine thiol-based transcriptional regulators orchestrate coordinated regulation of redox homeostasis and other cellular processes by “sensing” or detecting a specific redox-active molecule, which in turn activates the transcription of a specific detoxification pathway. The extent to which these sensors are truly specific in cells for a singular class of reactive small molecule stressors, e.g., reactive oxygen or sulfur species, is largely unknown. Here we report novel structural and mechanistic insights into a thiol-based transcriptional repressor SqrR, that reacts exclusively with organic and inorganic oxidized sulfur species, e.g., persulfides, to yield a unique tetrasulfide bridge that allosterically inhibits DNA operator-promoter binding. Evaluation of five crystallographic structures of SqrR in various derivatized states, coupled with the results of a mass spectrometry-based kinetic profiling strategy, suggest that persulfide selectivity is determined by structural frustration of the disulfide form. This energetic roadblock effectively decreases the reactivity toward major oxidants to kinetically favor formation of the tetrasulfide product. These findings lead to the identification of an uncharacterized repressor from the increasingly antibiotic-resistant bacterial pathogen, Acinetobacter baumannii, as a persulfide sensor, illustrating the predictive power of this work and potential applications to bacterial infectious disease.

---

Regulatory posttranslational modifications on thiol groups of redox-active and allosterically important cysteines are a critical component of signaling and redox regulation in cells^1–3^. The successful adaptation of microorganisms to an ever-changing microenvironment depends, to a considerable degree, on the specificity of the regulation of the expression of needed repair enzymes^4–6^. Understanding the extent to which transcriptional regulation is specific to a particular redox-active small molecule(s) is complicated by cross-talk or interconversion among these species in cells^7,8^ and the difficulties associated with identifying a posttranslational modification that is biologically meaningful at cell-appropriate concentrations that induce a cellular response^9^. Prominent exceptions to this are a handful of transcriptional regulators that employ a metallated cofactor, such as a mononuclear iron^10,11^, an iron–sulfur cluster^12,13^ or a heme-based sensing moeity^14^; here, the local inorganic chemistry alone is typically sufficient to explain the specificity of these switches toward their inducer^15^. In fact, it remains unclear if and how a non-metallated, dithiol-based transcriptional regulator achieves a similar level of specificity toward different cellular oxidants.

Recent work suggests that the upregulation of host-derived or bacterially-produced hydrogen sulfide (H_2_S) and downstream reactive sulfur species (RSS) is protective against a myriad of oxidative and nitrosative stressors in cells or animals infected by a number of medically important bacterial pathogens^4,16–20^. However, the cellular concentrations of H_2_S and more reactive RSS must be controlled to leverage these beneficial effects of H_2_S, while avoiding cellular toxicity^21^. We and others have discovered bacterial dithiol-based persulfide-sensing transcriptional repressors that are structurally unrelated and yet regulate the transcription of a subset of common downstream H_2_S detoxification genes^22–27^. Here, we focus our efforts on a model RSS-sensing, dithiol-based transcriptional regulator that is anticipated, on the basis of the enzymes encoded by regulated genes, to exhibit a high degree of specificity toward RSS^25^ and no meaningful response toward other non-sulfur based oxidants. Our model system is the sulfide-responsive transcriptional repressor, SqrR, that functions as a master regulator of sulfide-dependent gene expression in the purple photosynthetic bacterium *Rhodobacter capsulatus*^24^.

We exploit an MS-based, anaerobic assay to study the reactivity of SqrR toward a wide repertoire of redox active small molecules in a time-resolved manner, focusing our efforts on C9S SqrR which is active in cells and is functionally^24^ and structurally identical to wild-type SqrR, so as to minimize complications from oxidative chemistry occurring at non-conserved Cys9. In this assay we employ quantitative capping by excess iodoacetamide (IAM) in the absence of denaturing agents as confirmation that the protein is fully reduced (Figure S1A). Remarkably, we observe no change in this mass spectrum upon IAM capping following a 1 h incubation with a wide variety of oxidants including H_2_O_2_ and cysteine disulfide, CSSC (Figure 1A). In contrast, reduced SqrR readily reacts with inorganic (Na_2_S_4_) and organic (GSSH, SNAP persulfide donor^28^) persulfide donors shifting the mass distribution to that largely corresponding to a +62-Da species, a mass shift consistent with an intramolecular (intraprotomer) tetrasulfide crosslink between the conserved C41 and C107 residues (Figure 1B). Moreover, the tetrasulfide crosslink is stable over a wide range of pH (5-8) and cannot be capped with IAM; this suggests that the presumptive tetrasulfide is kinetically and thermodynamically stable and is not interconverting with a two-electron reduced persulfide moiety that might be present on each cysteine residue.

**FIGURE 1.**
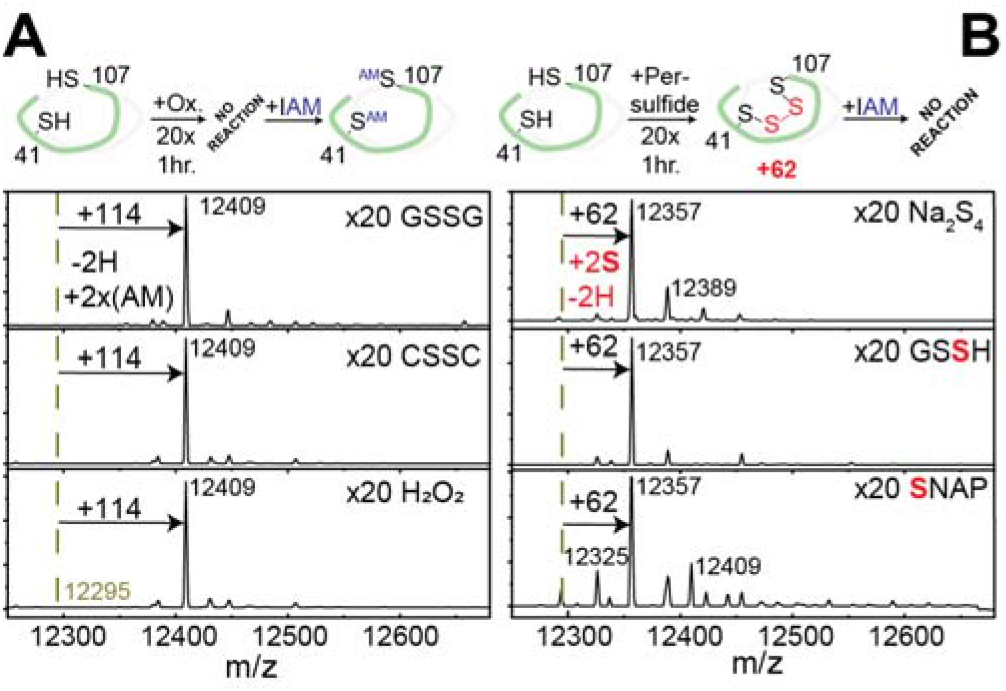
Specificity of C9S SqrR oxidation. C9S SqrR does not react with glutathione disulfide (GSSG), H_2_O_2_ or CSSC to form intramolecular crosslinks (A) and readily reacts with *in situ* generated glutathione persulfide (GSSH), Na_2_S_4_ and a SNAP persulfide donor (B).

These results show that the SqrR tetrasulfide crosslink is formed nearly exclusively in the presence of organic and inorganic persulfide donors and the more stable organic LMW thiol-derived disulfides do not efficiently react with C41 and/or C107. Moreover, we show that more stable precursors to oxidized sulfur species in cells, such as cysteine trisulfide, CSSSC^29,30^, do not readily react with C9S SqrR unless the reaction is done in the presence of residual (≈10%) cysteine monoxide, a potent electrophile (Supplementary information, Figure S2). Overall, these data suggest that other types intramolecular crosslinks might be disfavored in SqrR. In an effort to force C9S SqrR to form a disulfide bond, we took advantage of the fact that SqrR is able to react with strong electrophiles such as IAM and performed a reaction with TMAD (diamide), an electron deficient diazene commonly used to drive covalent modifications of thiols that ultimately lead to a disulfide bond in cells (Figure S1B). While the disulfide crosslinked form of SqrR is accessible under these conditions, it is unlikely it would be formed in in the cytoplasm, since it maintains a reduction potential consistent with a high GSH/GSSG ratio (≈ −220 mV)^31^.

We next measured the nucleophilicity of each Cys toward a neutral electrophile, *N*-ethylmaleimide (NEM), using a ratiometric pulsed alkylation-mass spectrometry approach with “heavy (*d*_5_-NEM)” and “light (H_5_-NEM)” variants of the reagent (Figure 2A). These experiments reveal that, as expected, C9 is highly reactive, with C107 and C41 less reactive (Figure S3). Examination of the kinetics and pH-dependence of the reactivity of each thiol in C9S SqrR and the corresponding single cysteine substitution mutants reveals that C41 is the less reactive Cys but shares with C107 an apparent p*K*_a_ of a typical solvent exposed thiolate (around 8) (Figure 2B, Figure S4). This relative stability of the protonated form of both thiolates partially explains the low reactivity towards oxidants, given the known correlation between p*K*_a_, redox potential and chemical reactivity^32,33^. This does not explain, however, why these less nucleophilic cysteines readily react with persulfides, which are weak electrophiles, to form a tetrasulfide bridge^34,35^.

**FIGURE 2.**
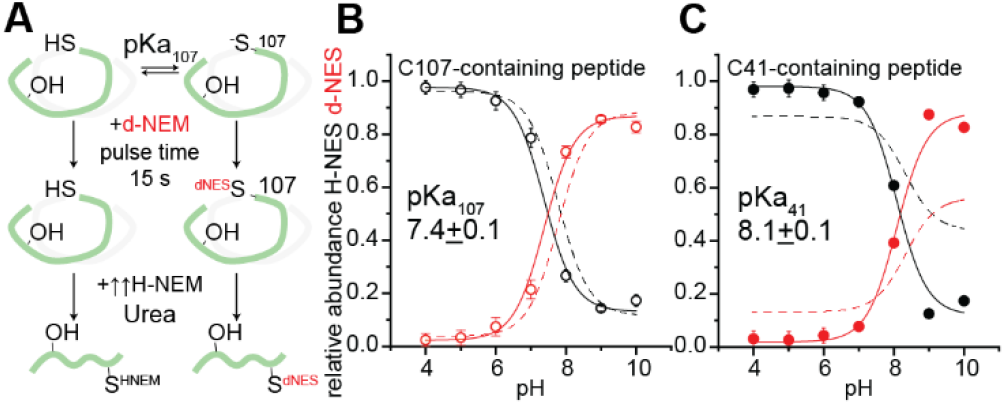
Apparent p*K*_a_ obtained from the pH dependence of the ratiometric pulsed alkylation-mass spectrometry analysis at 15s for each Cys residue in C9S/C41S SqR (A, B) and C9S/C107S SqrR (C). Dashed lines represent the fitting to the same peptide in C9S SqrR. See Figure S3 for C9S SqrR data, kinetic data and reaction schemes.

In order to determine the structural basis of persulfide selectivity and obtain further insights on the mechanism of tetrasulfide bond formation in SqrR, we first solved the structures of reduced wild type and C9S SqrRs to 2.12 Å and 1.36 Å (Supplemental table 1). These structures are remarkably similar (Figure S5) and reveal the expected arsenic repressor (ArsR) superfamily α1-α2-α3-α4-β1-β2-α5 family fold, with C41 and C107 derived from the α2 and α5 helices, respectively, and the S^γ^ atoms separated by 7.0 Å (Figure 3A-B). Both structures are missing a 17-residue N-terminal tail that is highly dynamic in solution (Figure S6), and that harbors C9, thus providing further validation of C9S SqrR as an excellent model for wild-type SqrR in chemical reactivity studies. C107 and C41 reside in a solvent accessible cavity at the top of the molecule formed by α2, α5 and α1’ helices (Figure 3A-B). Consistent with our pulsed alkylation experiments (Figure 2), C107 is more solvent-exposed (45% of the sidechain) while C41 in the cavity bottom is largely shielded from solvent (3% and 16% of the sidechain in C9S and WT SqrR, respectively). The overall electrostatics surface around each cysteine is also conserved with weak negative potential near the C-terminal α5 helix, while the interior of the cavity is characterized by significant positive potential (Figure 3).

**FIGURE 3.**
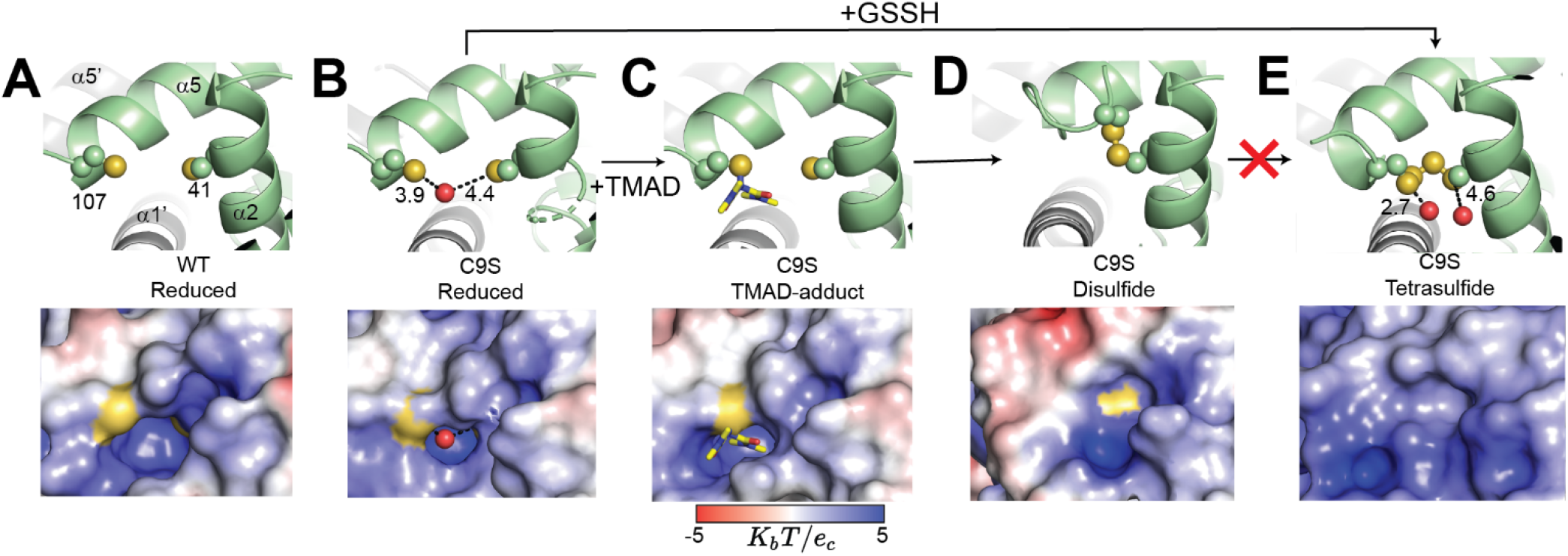
Thiol reactivity is determined by steric hindrance. Ribbon representation of protein structure in the proximity of the dithiol sensing site for reduced wild-type (WT) and C9S SqrRs in different oxidation states (top) with electrostatic surface renderings of the same region shown (bottom): a) reduced wild-type SqrR (pdb code 6O8L); b) reduced C9S SqrR (6O8K); c) TMAD soaked C9S SqrR (6O8N); d) TMAD-treated C9S SqrR (6O8O); e) GSSH-treated C9S SqrR (6O8M). *Yellow* spheres, sulfur atoms; *red* spheres, H_2_O oxygen atoms. See Supplement table 1 for structure and refinement statistics.

Remarkably, soaking the reduced C9S SqrR crystal with TMAD results in the formation of what is best described as an on-pathway intermediate to disulfide bond formation, and features what is to our knowledge the first structure of sulfenamide linkage with a diazene compound (solved at 1.46 Å resolution). One of the diazene nitrogen atoms of TMAD forms an S-N bond with the more nucleophilic C107 S^γ^ atom, blocking the access to the cavity where C41 remains reduced (Figure 3C). This structure suggests that modification of C107 at the edge of the cavity creates a steep energetic barrier to both subsequent nucleophilic attack on that sulfonamide S by the C41 S^γ^ that would lead to a disulfide bond, or nucleophilic attack of C41 S^γ^ on a low molecular weight electrophile. Indeed, the crystallographic structure of the disulfide-crosslinked SqrR (Figure 3D) reveals a high energetic barrier in the reaction coordinate to form the disulfide, given the large structural rearrangement that must occur for the disulfide to form. An analysis of structural frustration (Figure S7) suggests that formation of the disulfide conflicts with folding in the crystal structure, characterized by very high local frustration when compared to the reduced state (Figure 4). These highly frustrated contacts derive from the unfolding of one turn of the C-terminal α5 helix around C107, accompanied by increased solvent exposure of F106 and Y103 and a decreased solvent accessibility of the disulfide bond.

**FIGURE 4.**
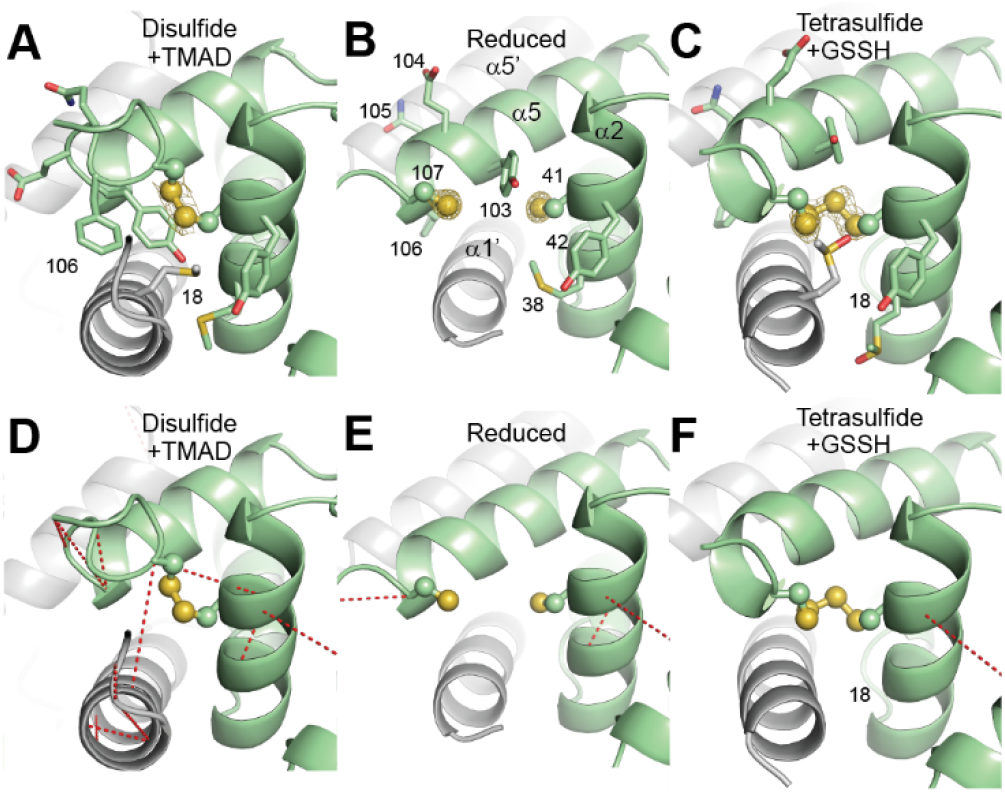
The disulfide crosslink introduces high local structural frustration relative to the tetrasulfide form. Protein structure in the proximity of the dithiol site for C9S SqrR in the disulfide (A), reduced (B) and tetrasulfide (C) states are shown with the electron density corresponding to the cysteine sulfur atoms, highly frustrated contacts (*red* broken lines) are shown for each structure (D-F).

While the disulfide and the TMAD-adduct structures argue that the lack of reactivity toward reactive oxygen- and disulfide based-oxidants could also be imposed by the required local unfolding as well as the stability of the protonated thiolate, they do not provide a molecular rationale for formation of a dominant tetrasulfide crosslinked species. To investigate this, we solved the structure of the tetrasulfide species of C9S SqrR, obtained by reaction with an *in situ* organic persulfide. While this structure (Figure 3E, Figure 4C) reveals a covalent, tetrasulfide bond between C41 and C107 as expected from the mass spectrometry, the tetrasulfide species, like the disulfide state, features collapse of the dithiol pocket, completely shielding the tetrasulfide from solvent, particularly so for the two additional GSSH-derived sulfur atoms (Figure 3E). This low solvent accessibility likely strongly attenuates the reactivity of these bridging sulfur atoms toward strong electrophiles, *e.g*., iodoacetamide (Figure 1), and other nucleophiles, *e.g.*, LMW thiols that are generated *in situ* during the reaction, at least at these concentrations. Moreover, this form is well folded with a decrease of highly frustrated contacts in the cysteine pocket, suggesting that the local contacts within that pocket stabilize the folded state even more so the reduced state (Figure 4F).

It is interesting to note that the disulfide- and tetrasulfide-crosslinked SqrR species each show significant local structural perturbations only in the immediate vicinity of C107 and the α 5 helix, while the DNA-binding helices and indeed the global structure of the dimer remain largely unaffected (Figure S5). However, the protein DNA-binding affinity is markedly impacted by these covalent modifications, with an allosteric coupling free energy (Δ*G*_c_) ≈4.5 kcal·mol^−1^ for tetrasulfide crosslinked SqrR and a somewhat smaller value for the disulfide form (Supplement table 2 and Figure S8). This characteristic absence of a large global conformational change upon inducer recognition is an emerging feature of ArsR family repressors^36,37^.

While these structures hint at the mechanism of disulfide bond formation by strong electrophiles like TMAD, they do not provide direct insights into the mechanism of tetrasulfide bond formation. To this end, we conducted a series of kinetic reactivity experiments with various organic persulfide donors followed by capping with excess IAM at variable incubation times (Figure 5A). The conversion of the reduced form and formation of the tetrasulfide bridge are kinetically well-modelled with an identical pseudo first order rate constant (Figure 5B). The minor di- and trisulfide species detected in these profiles are, on the other hand, kinetically well-modeled as parallel reactions derived from the same reduced SqrR, rather than as on-pathway intermediates, with much slower pseudo-first order reaction rates independent of the identity of the sulfur donor (Supplemental table 3). A single end-point reactivity assay of the disulfide-crosslinked C9S SqrR with GSSH indeed confirms that the disulfide is not on-pathway to the tetrasulfide product (Figure S9). The rate constant of tetrasulfide formation can be modulated by the concentration and identity of the persulfide donor, the presence of bound DNA or heme-based (R)S• radical generators^38^ (Supplemental table 3). By increasing the donor concentration or carrying out reactions in the presence of hemin, we could observe significant accumulation of an on-pathway persulfide intermediate and fit each dataset with a sequential reaction mechanism (Figure 5C, Figure S10). This persulfide intermediate is likely formed on C107, since it is the more nucleophilic cysteine (Figure 5D). We hypothesize that this initial persulfidation is followed by subsequent reactions on one or both Cys where GSSH/GSS^−^ can ultimately drive the formation of the tetrasulfide by exploiting the dual character of persulfides and their increased nucleophilicity when compared to the thiols (Figure 5D). Single time point GSSH experiments with single Cys-containing mutants confirm that mixed disulfides and polysulfides with glutathione can potentially be formed on either Cys (Figure S9). However, given the steric hindrance and reduced reactivity of the C41 thiol, these subsequent reactions with the persulfide donor likely occur more rapidly on C107; in this case, modification of C41 would occur by intramolecular sulfur donation (Figure 5D).

**FIGURE 5.**
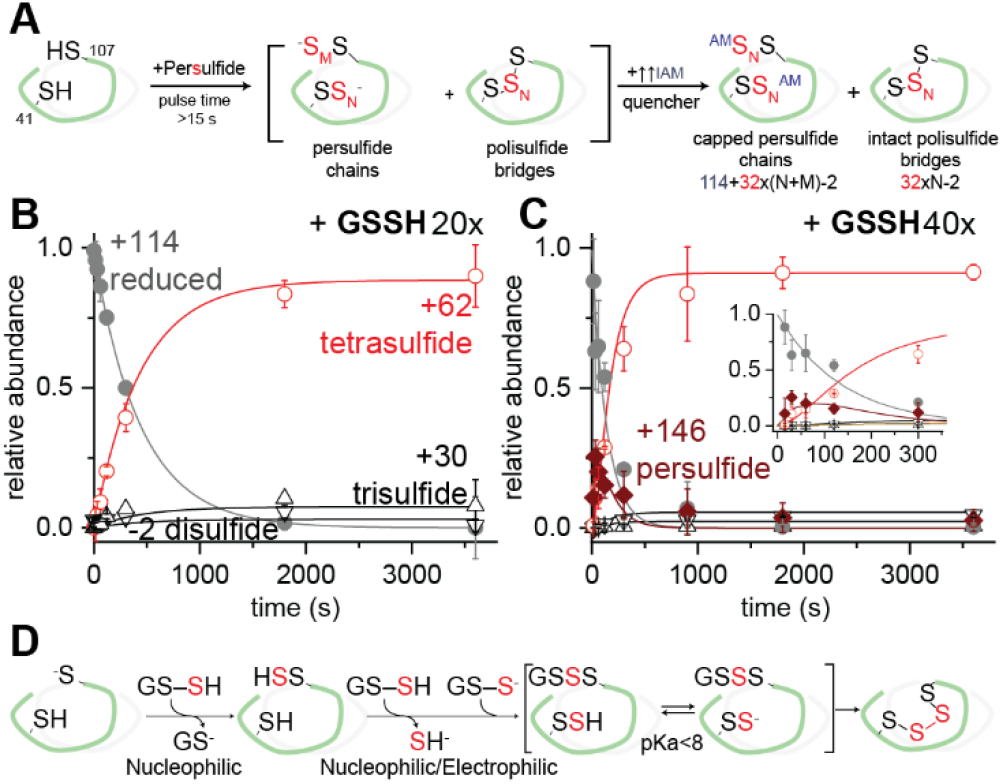
The kinetics of tetrasulfide bond formation in SqrR. (A) Schematic representation of the pulsed-chase approach to study the kinetics of the reaction with RSS and the expected outcome. (B, C) Relative abundance of the indicated species as a function of pulse time. Continuous liines in panel B and C represent fits to kinetic mechansims 1 and 2 (Supporting Information), respectively. (D) Mechanistic proposal compatible with the results presented.

Our results define three key determinants of persulfide selectivity in dithiol based transcriptional regulators: 1) a major energetic barrier to formation of the disulfide attributed to local structural frustration that allows kinetic partitioning to the tetrasulfide product; 2) moderately basic thiols that are poorer nucleophiles at physiological pH, and 3) a decrease in the reactivity of the less nucleophilic thiol upon formation of the initial intermediate (persulfide, mixed disulfide or S-N adduct). These observations make the prediction that other dithiol-based transcriptional regulators that share a similar molecular architecture might share similar chemical reactivity characteristics. To test this idea, we purified and characterized an as yet uncharacterized dithiol-based repressor encoded by *Acinetobacter baumannii*, denoted *Ab* BigR (37% sequence identity), and show that this sensor, like *Rc* SqrR (Figure 6), reacts selectively with GSSH over other oxidants, albeit with somewhat slower kinetics and incorporation of an extra sulfur atom in the major product (pentasulfide adduct). This is consistent with a minimal structural perturbation of a somewhat larger cavity as inferred from a structure of the homologous protein with putative RSS selectivity from *Xylella fastidiosa* (43% identity, Figure S11)^39^. Consistent with the physical properties presented here, *Ab* BigR has been recently shown to regulate a three-gene operon encoding a classic persulfide dioxygenease (PDO) and two poorly characterized membrane proteins involved in sulfur transport (YedE/YeeE), confirming a cellular role in RSS sensing and H_2_S homeostasis^40,41^.

**FIGURE 6.**
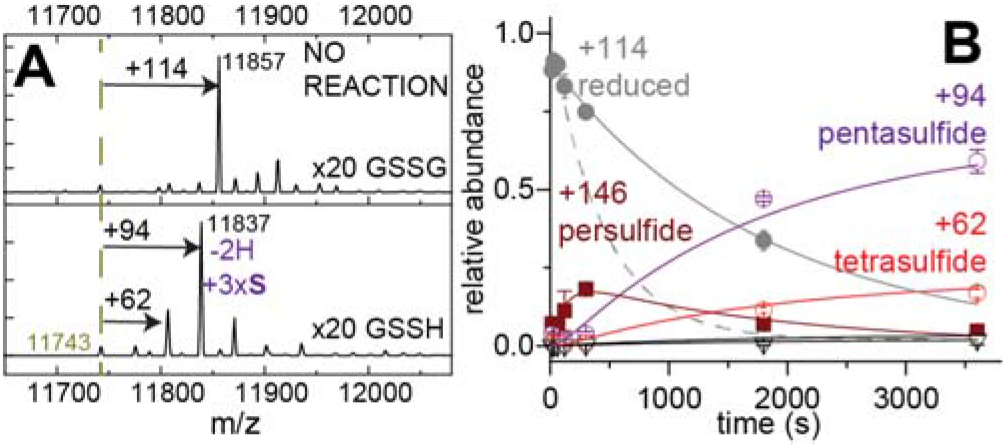
*Ab* BigR reactivity follows a SqrR pattern with a longer polysulfide chain (A) and slower kinetics (B).

In conclusion, we demonstrate that it is indeed possible for non-metallated, dithiol-based transcriptional regulators to achieve a level of specificity required for bacterial cells to trigger a unique response to RSS/H_2_S toxicity rather than general redox misbalance. This is functionally important since increased H_2_S biogenesis has been shown to be necessary for bacteria to counteract the effect of antibiotic stress and host derived reactive oxygen and nitrogen species stressors^4,16,19,42^. We show that this chemical specificity derives from a unique energetic landscape dictated by the protein matrix, which dictates the spectrum of posttranslational modifications by minimizing local structural frustration, while exploiting steric hindrance and low nucleophilicity. This work illustrates how the “minimal frustration principle” previously applied to allosteric communication^43,44^ and enzymatic activity^45^ dictates chemical specificity when supramolecular chemistry determined by the protein structure is critical to understand biological outcomes.

## MATERIALS AND METHODS

### Protein Preparation

Overexpression plasmids encoding C9S, C9S/C41S and C9S/C107S SqrRs were constructed by PCR-based site-directed mutagenesis using pSUMO-CzrA as template^24^ and verified using DNA sequencing. Plasmids used for the expression of wild-type SqrR were reported previously^24^. Proteins were expressed in *E. coli* BL21(DE3)/pLysS cells and purified as previously described for SqrR^24^.

*Ab*BigR was subcloned into a pHis plasmid between *Nco*I and *Ned*I sites, allowing downstream removal of the N-terminal His_6_ tag. *Ab*BigR was expressed in *E. coli* BL21(DE3)/pLysS cells and purified using a variation of a previously described protocol^46^. Briefly, freshly harvested cell expressing *Ab*BigR paste was suspended in 200 mL of Buffer B (25 mM MES, 750 mM NaCl, 2 mM TCEP, 1 mM EDTA, pH 6.0) and lysed by sonication using a Fisher Scientific model 550 sonic dismembrator. The cellular lysate was centrifuged at 8000 rpm for 15 min at 4 °C. The supernatant was collected and subjected to protein and nucleic acid precipitation using 10% polyethylenimine (0.015 v/v) at pH 6.0. After stirring for 1 h at 4 °C, the solution was centrifuged at 8000 rpm for 15 min at 4 °C. The supernatant was precipitated by the addition of (NH_4_)_2_SO_4_ to 70% saturation with stirring for 2 h. After centrifugation at 8000 rpm for 15 min, the precipitated protein was dissolved and dialyzed against Buffer A (25 mM MES, 75 mM NaCl, 2 mM TCEP, 1 mM EDTA, pH 6.0). This solution was loaded onto a 10 mL SP (sulfopropyl) Fast Flow cation exchange column equilibrated with Buffer A. The protein was then eluted using a 150 mL linear gradient of 0.075-0.75 M NaCl.

C9S SqrR samples for backbone assignments were isotopically labeled using published procedures^47^ with all isotopes for NMR experiments purchased from Cambridge Isotope Laboratories. Selenomethionine labelled C9S SqrR was obtained using a published procedure^48^.

All of the proteins characterized here eluted as homodimers as determined by a calibrated Superdex 75 (GE Healthcare) gel filtration chromatography column (25 mM Tris, 0.2 M NaCl, 2 mM EDTA, 2 mM TCEP, 5% glycerol, pH 8.0, 25 °C).

### X-ray Crystallography

All proteins were crystallized at 20 °C by a hanging-drop vapor diffusion method at a concentration of 6 mg/mL in the crystallization buffer (25 mM Tris, 0.2 M NaCl, 2 mM EDTA, 5% glycerol, pH 8.0). WT SqrR was crystallized in the presence of 2 mM TCEP with a mother *liquor* containing 2.5 M ammonium sulfate, and 0.1 M sodium acetate pH 4.6. Reduced C9S SqrR was crystallized in the presence of 2 mM TCEP with a mother *liquor* containing 1.8 M sodium citrate, pH 6.4. We could distinguish no difference in the electron density maps when TCEP was omitted in C9S SqrR crystallization trials. To obtain a structure of C9S SqrR diamide intermediate, 1 mM diamide was added into the droplets containing crystals of C9S SqrR in the absence of TCEP and the soaked crystals were collected after 1 h.

C9S SqrR in the tetrasulfide state was obtained following a 1 h incubation with GSSH at 20-fold molar excess in 150 mM phosphate buffer pH 7.4 and the completion of the reaction was monitored by LC-ESI-MS. The protein was buffer exchanged in anerobic conditions into the crystallization buffer. Tetrasulfide crosslinked C9S SqrR was crystallized with a mother liquor containing 0.2 M ammonium sulfate, 0.1 M sodium cacodylate, pH 6.5 and 30% PEG 800.

C9S SqrR in the disulfide state was obtained by incubation of SeMet-labelled C9S SqrR for 10 min anaerobically in a desalting column in a solution containing 10 mM TMAD and the completion of the reaction was monitored by LC-ESI-MS. The protein was buffer exchanged in anerobic conditions into the crystallization buffer. Disulfide C9S SqrR was crystallized with a mother liquor containing 0.1 M Bis-tris propane. pH 5.5-6.5 and 1.6-1.85 M ammonium sulfate.

For cryoprotection, crystals were transferred for a few seconds into a reservoir solution supplemented with 20% (v/v) glycerol and were subsequently flash-frozen in liquid nitrogen. Diffraction data were collected at 100 K at the 4.2.2 beamline at the Advanced Light Source (Berkeley, CA). The data were indexed, integrated, and scaled using the XDS package.

The structure of reduced C9S SqrR form (PDB code 6O8L) was solved by molecular replacement using PHASER and the PDB code 3PQJ as search model^49^. Phase calculations were performed using Phaser in PHENIX AutoMR module^50^. The C9S SqrR disulfide state (PDB code 6O8O) form was phased using Single Anomalous Diffraction (SAD) methods of a selenomethionine derivative protein. The rest of variants were phased by molecular replacement using the reduced form of C9S as the search model. In all cases, the Autobuild function was used to generate an initial model that was improved by iterative cycles of manual building in Coot^51^ and refinement using PHENIX^50^. MolProbity software^52^ was used to assess the geometric quality of the structural models and Pymol (http://www.pymol.org) was used to generate molecular images. The electron densities of sulfur atoms are indicated in Figure 4 for the disulfide, reduced and tetrasulfide states, while the SqrR diamide intermediate is only shown in Figure 3 without the electron density for clarity. The authors note that the electron density of the S-N is very well define while the carbonyl on the other N is likely in more than one conformation.

#### Data deposition

Table S1 provides the statistics for data collection, processing, refinement and the filenames deposited to the PDB (PDB ID: 6O8L, 6O8K, 6O8N, 6O8O, 6O8M).

The conformational frustration patterns were calculated using the protein Frustratometer software (www.frustratometer.tk/)^45,53^. This software that has been described in more detail elsewhere^45,53^ and has been extensively used to classify energetic frustration in different folded proteins^44,45,53,54^. It determines the local frustration by evaluating how favorable a specific contact is relative to the set of all possible contacts in that location. Here we have focused the analysis on the configurational frustration as a mean for comparing the folded structures of the different oxidation states of SqrR. This way of measuring frustration imagines that the residues are displaced in location. The energy variance thus reflects contributions to different energies of other compact conformations. In this calculation the decoy energy set involves randomizing the identities, the distance and densities of the interacting amino acids. A contact is defined as ‘minimally frustrated’ if its native energy is at the lower end of the distribution of decoy energies (*i.e.*, most of the other distances in that position would be unfavorable). Conversely, a contact is defined as 'highly frustrated' if its native energy is at the other end of the distribution (*i.e.*, most of the other distances for that amino acid pairs in that position would be more favorable for folding than the native ones by more than one standard deviation of that distribution). If the native energy is in between these limits, the contact is defined as ‘neutral’.

In Figure 4 (main text), minimally frustrated and neutral contacts are omitted to focus in the increase of the highly frustrated contacts in the disulfide state when compared to the reduced and tetrasulfide state. All the minimally frustrated contacts can be found in Figure S7, with only the neutral contacts omitted for clarity.

### Preparation of Glutathione Persulfide

Glutathione persulfide (GSSH) was freshly prepared by mixing a five-fold molar excess of freshly dissolved Na_2_S with glutathione disulfide and incubating anaerobically at 30 °C for 30 min in degassed 300 mM sodium phosphate (pH 7.4). The concentration of the *in situ* generated GSSH was determined using a cold cyanolysis assay^55^ and used without further purification at the indicated final concentrations.

### Synthesis of cysteine trisulfide

Cysteine trisulfide (CSSSC) was prepared with minor modifications from that previously reported^29,56^. To a solution of 1 g of cystine in 10 mL 1 M H_2_SO_4_ in an ice bath, 630 mg of peracetic acid was added dropwise and the reaction was stirred on ice overnight. The pH was adjusted to 4 with cold pyridine and cystine-S-monoxide intermediate was precipitated with 3X volume cold ethanol and collected by vacuum filtration. The intermediate was then dissolved in 10 mL 1 M H_2_SO_4_ to which 650 mg of Na_2_S was added and the reaction was sealed and stirred at room temperature for 1 hour. The pH was then adjusted to 3.5 using pyridine and cysteine trisulfide was precipitated by addition of 3X volume of a 50/50 mixture of ethanol/THF. The precipitate was filtered, washed with 50/50 ethanol/THF and dried. Mass spectral analysis (ESI-MS) *m/z* [M + Na]^+^ calculated for C_6_H_12_N_2_O_4_S_3_Na is 295.35 (expected); found 295.20 (observed).

### Ratiometric Pulsed-Alkylation Mass Spectrometry (rPA-MS)

#### Alkylation step

Sample preparation was adapted from earlier reports^57,58^ and optimized for C9S SqrR. All experiments ware performed anaerobically in a glovebox. The purified C9S SqrR was first buffer exchanged into buffer C (25 mM HEPES pH 7.0, 200 mM NaCl, 5 mM EDTA) to remove the reducing reagent. The pulsed alkylation reagent *d*5-*N*-ethylmaleimide (*d*5-NEM, Isotech, Inc.) was dissolved in acetonitrile at 25 mM. C9S SqrR (monomer concentration of 50 μM) was allowed to react with a 3-fold thiol excess of *d*5-NEM. At different pulsed time intervals (*t*), a 50 μL aliquot was removed and added to 50 μL of chase solution containing a 900-fold thiol excess of protiated *N*-ethylmaleimide (H5-NEM), 100 mM Tris-HCl, pH 8.0, and 8 M urea. After a 1 h chase, the quenched reaction was removed from the anaerobic chamber and precipitated by adding a final concentration of 12.5% trichloroacetic acid (TCA) on ice for 1.5 h. The precipitated protein pellets were collected by centrifugation at 4 °C for 30 min, with the supernatant discarded and the pellet washed by iced acetone three times. After the final wash, the pellet was dried by speed-vacuum centrifugation at 45 °C for 10 min. The same experiments were performed with the C9S/C107S SqrR and C9S/C41S SqrR. For single time point pulsed-alkylation at different pH, buffers with a range of pH were prepared (100 mM citrate, 200 mM NaCl, 2mM EDTA, pH 4.0, pH 5.0, pH 6.0; 100 mM HEPES, 200 mM NaCl, 2mM EDTA, pH 7.0, pH 8.0; 100 mM Tris-HCl, 200 mM NaCl, 2 mM EDTA, pH 9.0; 100 mM CAPS, 200 mM NaCl, 2 mM EDTA, pH 10.0). C9S SqrR and its mutant (C41S/C107S) (monomer concentration of 50 μM) was mixed with a 3-fold thiol excess of *d*5-NEM at the different pH conditions. At a 15 s pulse time, a 50 μL chase solution containing a 900-fold thiol excess of protiated *N*-ethylmaleimide (H5-NEM), 100 mM Tris-HCl, pH 8.0, and 8 M urea, was added to 50 μL of pulsed solution to quench the alkylation. Same sample washing protocol as mentioned above were used to prepare the single pulse time sample for protease digestion.

#### Protease digestion and MALDI-TOF Mass Spectrometry

Lys-C protease was used to produce the dominant peptides containing either [C107] or [C41], while trypsin was used to obtain the dominant peptides containing [C9]. For digestion, the vacuum dried protein pellets were re-suspended in 20 μL digestion buffer (20 mM ammonium bicarbonate, 10% acetonitrile, 2 M Urea, pH 8.0) with 100 ng of the protease added. Both protease digestions were performed at 37 °C for 20 h. Protease digestions were quenched by the addition of 2 μL 10% trifluoroacetic acid (TFA) and the digestion products first purified by C-18 Zip-tip before spotting on MALDI-plate with ◻-cyano-4-hydroxycinamic acid (CCA) matrix at a 5:1 matrix:sample ratio for mass spectrometry analysis. The MALDI-TOF mass spectra data of all the samples were collected by using a Bruker Autoflex III MALDI-TOP mass spectrometer with 200 Hz frequency-tripled Nd:YAG laser (335 nM).

#### MALDI-TOF data analysis

In the MALDI-TOF spectra of the Cys-containing Lys-C and trypsin generated peptides, the relative abundances of the *d*5-NEM and H5-NEM modified 85-110 (TIYYSLSDPRAARVVQTVYEQF**C**SGD), 30-48 (ALAHEGRLMIM**C**YLASGEK) and 9-20 (**C**AALDAEEMATR) peptides obtained from the SqrR variants were quantified averaging the intensity of the [M+H]^+^ most abundant isotopomers. The kinetics were fitted to a single or double exponential decay since these reactions are expected to follow a pseudo first-order kinetics (as indicated in Figure S4). The kinetics constants obtained are as expected for a moderately nucleophilic cysteine at basic pH values, while they behave as protected or weakly nucleophilic cysteines at acidic pH values^57,58^.

### LC-ESI-MS Analysis of Derivatized Proteins at Different Pulse Times

Reduced C9S SqrR and BigR were buffer exchanged anaerobically into 150 mM potassium phosphate buffer pH 7.4 containing 1 mM EDTA. Reactions containing 60 μM protein were anaerobically incubated at room temperature with a 20-fold excess of oxidizing reagent, *e.g*., GSSH for 1 h or for different times as indicated in the figures. The reaction was quenched by addition of equal volume 60 mM iodoacetamide (IAM). The reactions were then washed 3 times with 150 mM potassium phosphate buffer pH 7.4 containing 1 mM EDTA to remove excess reagents and analysis was performed in the Laboratory for Biological Mass Spectrometry at Indiana University using a Waters Synapt G2S mass spectrometer coupled with a Waters ACQUITY UPLC iClass system. 5 μL protein samples were loaded onto a self-packed C4 reversed-phased column and chromatographed using an acetonitrile-based gradient (solvent A: 0% acetonitrile, 0.1% formic acid; solvent B: 100% acetonitrile, 0.1% formic acid). Data were collected and analyzed using MassLynx software (Waters).

The relative intensities of each species were obtained from the convoluted intensities of peaks corresponding to the [P+8H]^8+^, [P+9H]^9+^, [P+10H]^10+^, [P+11H]^11+^, [P+12H]^12+^, [P+13H]^13+^, [P+14H]^14+^, and [P+15H]^15+^ charge states, assuming that different oxidation states had identical efficiencies. Only the species with relative intensities higher than 10% in at least one time point were included in the analysis, except for disulfide and trisulfide that were included and fitted to evaluate them as intermediates or parallel reaction products.

The kinetic data were fitted with Dynafit^59^ following a minimal kinetic model pseudo-order in the persulfide donor. The order for GSSH was estimated from the pseudo first order constant for the formation of the tetrasulfide. The intermediate (persulfide) were fitted to a simple consecutive reaction model, while the two minor species (disulfide and trisulfide) were fitted to simple parallel reaction. Two kinetic models were used to fit the data:

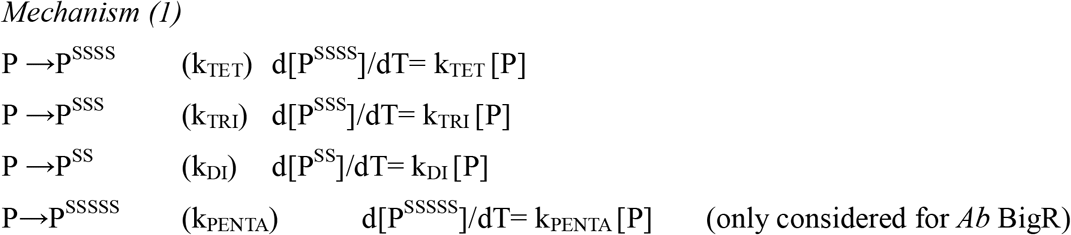

where P represents the reduced protein monomer (quantified from the relative intensity of the peak at +114 *m/z* IAM capped protein), P^SSSS^ represents the tetrasulfide bridge within the protein monomer monomer (quantified from the relative intensity of the peak +62 *m/z*), P^SSSS^ represents the trisulfide bridge within the protein monomer (quantified from the relative intensity of the peak +30 *m/z*), P^SS^ represents the disulfide bridge within the protein monomer (quantified from the relative intensity of the peak –2 *m/z*), P^SSSSS^ represents the penta sulfide bridge within the protein monomer (quantified from the relative intensity of the peak +94 *m/z* considered only for *Ab* BigR), k_TET_ is rate constant for formation of the tetrasulfide, k_TRI_ is rate constant for formation of the trisulfide, k_DI_ is rate constant for formation of the disulfide and k_PENTA_ is rate constant for formation of the pentasulfide.

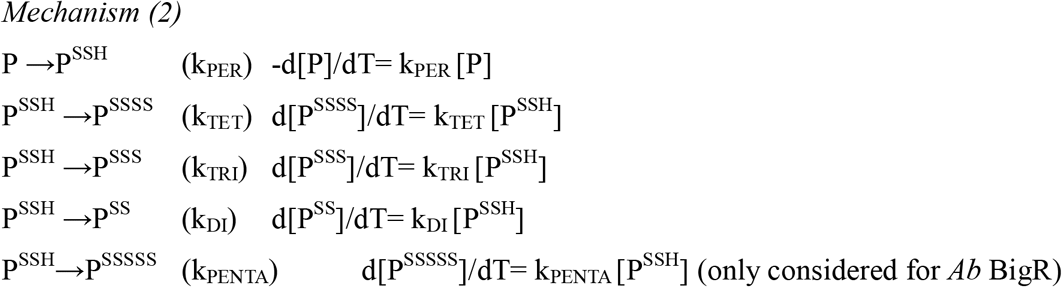

where the representation is the same as in Mechanism (1), but also including P^SSH^ as the representation of the persulfide intermediate protein monomer (quantified from the relative intensity of the peak at +146 *m/z* IAM capped species) and k_PER_ is rate constant for consumption of the P to form such persulfide intermediate.

No significant improvement of the fitting could be obtained by including reverse reactions or other parallel reactions.

These models and the obtained parameters imply that the rate limiting step is the formation of a persulfide on SqrR which is dependent on the concentration of persulfide donor. The slower reaction with the SNAP donor suggests that the rate of the reaction depends also on the nature of the persulfide donor.

The presence of persulfide radicals (as one would expect in the presence of hemin^38,60^) results in a two-fold increase in the kinetics of disappearance of reduced SqrR and corresponding production of the tetrasulfide. Hemin also gives rise to an accumulation of the persulfide intermediate as is observed at higher GSSH concentration. In contrast, oxidized (Fe^3+^) cystochrome c (Cyt^3+^) does not impact the persulfidation kinetics, as one would expect for a coordinately saturated (protected) heme group under the high phosphate concentrations and pH 7.4 used here (pH 7.4 is two pH units below the alkaline transition). Despite the fact that Cyt^3+^ does not substantively impact the kinetics of GSSH-dependent oxidation of SqrR, we do observe a change in the oxidation state of Cyt to Fe^2+^ in the presence of persulfides consistent with what has been previously reported^38^.

The parameters obtained for the reaction in the presence of DNA suggest that there is kinetic allosteric inhibition, *i.e*., allokairy^61^ for the reaction with GSSH which is as expected when one takes into account the negative thermodynamic linkage in this system (Supplementary table 2).

The reaction with BigR is significantly slower which may derive from a restriction imposed by a well-folded N-terminal helix that is significantly more than that attributed to the N-terminal disordered region (Figure S6) in SqrR.

### Fluorescence anisotropy-based DNA binding experiments

Standard fluorescence anisotropy-based DNA binding experiments were also carried out using two 29-bp fluorescein (F) labeled sqr operators DNA fragment, 5’F- (T)TGACAT**ATTC**ACAACTTCG**GAAT**GTAA-3’ and its complement (the ATTC-x_8_-GAAT *sqr* box is bold) from the rcc1451 promotor region (labeled 1451) and 5’F- (T)TGCCT**ATTC**ATATTCTC**ATAT**GTGC-3’ and its complement from the SQR promotor (labeled 0785) in the DNA binding buffer (10 mM HEPES, pH 7.0, 0.2 M NaCl, 1 mM EDTA in presence or in absence of 2 mM TCEP) exactly as described in previous work.^36^ After the titration was completed the protein was either dissociated from the DNA by addition of a 900-fold excess of IAM (when performed in the absence of reducing agent) or fully bound to the DNA by the addition of a quantitative amount of TCEP to either the tetra sulfide form. This experiment validated that the tetrasulfide formation is reversible and inhibitory of association with the DNA. All anisotropy-based data were fit to a simple 1:1, non-dissociable dimer binding model to estimate *K*_a_ using DynaFit.^59^

## ASSOCIATED CONTENT

### Supporting Information

Tables of kinetic parameters, crystallographic structure data collection and refinement statistics, DNA-binding constants; figures with details from crystal structures and primary experimental mass spectrometry data, and DNA-binding data.

## AUTHOR INFORMATION

### Author Contributions

The manuscript was written through contributions of all authors. All authors have given approval to the final version of the manuscript.

### Notes

The authors declare no competing financial interest.

## ACKNOWLEDGMENT

We thank Hongwei Wu with his help with NMR data acquisition. We thank Yu-Chen Huang for her help in protein purification. We thank Steve Xu and Ming Xian from Washington State University who kindly provided the SNAP donor. The authors gratefully acknowledge use of the Macromolecular Crystallography Facility at the Molecular and Cellular Biochemistry Department, Indiana University Bloomington. We also thank Jay Nix for his assistance during X-ray data collection at beamline 4.2.2, ALS.

## ABBREVIATIONS

RSS: reactive sulfur species
SqrR: sulfide quinone reductase repressor
LMW: low molecular weight
TMAD: tetramethylazodicarboxamide

