## Supporting Information for "Structural determinants of persulfide-sensing specificity in a dithiol-based transcriptional regulator"

**This file contains Supporting Tables S1-S3 and Supporting Figures S1-S11.**

**Supporting Information**

page

**Supporting Tables**


**Table S1.** Crystallographic structure data collection and refinement statistics S3

**Table S2.** Thermodynamic parameters for DNA binding with two cognate operators in SqrR variants S4

**Table S3.** Kinetic parameters for reactions with persulfide donors for SqrR and BigR S5


**Supporting Figures**


**Figure S1.** Illustration of quantitative capping of reduced C9S SqrR with IAM under native conditions and the reaction of C9S SqrR with TMAD. S6

**Figure S2.** Cysteine trisulfide synthesis and reaction with C9S SqrR. S7

**Figure S3.** rPA-MS single time point assays used to compare the relative reactivies of C9, C41 and C107 in SqrR variants. S8

**Figure S4.** rPA-MS kinetics and estimation of the p$K_a$ of C41 and C107. S9

**Figure S5.** $C^{\alpha}$ ribbon diagram superpositions and pair-wise $C^{\alpha}$ root-mean square deviations (RMSD) in pairs of SqrR structures solved here. S10

**Figure S6.** $^{1}H$, $^{15}N$ HSQC spectra and $^{1}H$–{$^{15}N$} heteronuclear nuclear Overhauser enhancement (hNOE) of reduced C9S SqrR. S11

**Figure S7.** Frustration patterns in C9S SqrR in various allosteric states. S12

**Figure S8.** SqrO–DNA binding isotherms for C9S and wild-type SqrRs in the reduced, disulfide and tetrasulfide crosslinked states. S13

**Figure S9.** The reactivity of the putative persulfide intermediate in single Cys-containing SqrR variants and kinetic trapping of the disulfide. S14

**Figure S10.** Representative kinetic traces that illustrate the time-course of reactivity of C9S SqrR toward various oxidants under various conditions. S15

**Figure S11.** Comparison of the structures and polysulfide product distributions of *Rc* SqrR and BigR-like RSS sensors. S16

**References** S17

### Supporting Tables

**Supplemental table 1.** Crystallographic structure data collection and refinement statistics

| | SqrR C9S | SqrR WT | SqrR C9S tetrasulfide | SqrR C9S disulfide | Sqr C9S - TMAD |
|---|---|---|---|---|---|
| *Data collection* | | | | | |
| Wavelength (Å) | 1.072152 | 1.072152 | 1.000031 | 0.97625 | 1.000031 |
| Space group | P6122 | P6122 | P4212 | P3221 | P6122 |
| *Cell dimensions* | | | | | |
| a, b, c (Å) | 46.38 46.38 159.64 | 46.85 46.85 163.31 | 84.84 84.84 57.26 | 92.60 92.60 148.98 | 46.51 46.51 158.47 |
| α, β, γ (°) | 90.00 90.00 120.00 | 90.00 90.00 120.00 | 90.00 90.00 90.00 | 90.00 90.00 120.00 | 90.00 90.00 120.00 |
| Resolution (Å) | 53.21 – 1.36 | 54.44 – 2.12 | 59.99 – 1.95 | 49.66 – 2.50 | 52.82 – 1.46 |
| $R_{sym}$ | 0.038 (0.858) | 0.081 (0.972) | 0.167 (1.147) | 0.112 (1.649) | 0.050 (0.944) |
| $R_{meas}$ | 0.039 (0.911) | 0.083 (0.997) | 0.174 (1.241) | 0.115 (1.729) | 0.051 (0.984) |
| $R_{pim}$ | 0.009 (0.297) | 0.019 (0.219) | 0.047 (0.340) | 0.025 (0.622) | 0.012 (0.273) |
| CC1/2 | 1.000 (0.757) | 1.000 (0.894) | 0.998 (0.790) | 1.000 (0.660) | 1.000 (0.875) |
| I/σ(I) | 44.3 (2.0) | 26.6 (3.7) | 14.7 (2.3) | 27.2 (1.7) | 33.9 (2.4) |
| Completeness (%) | 97.3 (79.9) | 100.0 (100.0) | 100.0 (100.0) | 99.7 (100.0) | 99.9 (100.0) |
| Multiplicity | 17.4 (8.7) | 19.4 (20.2) | 13.6 (13.0) | 21.1 (18.7) | 18.4 (12.5) |
| *Refinement* | | | | | |
| Resolution (Å) | 26.61 – 1.36 (1.41 – 1.36) | 27.22 – 2.12 (2.19 – 2.12) | 42.42 - 1.95 (2.02 – 1.95) | 49.66 – 2.50 (2.59 – 2.50) | 40.29 – 1.46 (1.55 – 1.46) |
| No. unique reflections | 22169 | 6617 | 14748 | 24511 | 17998 |
| $R_{work}$ | 0.1480 | 0.2233 | 0.2083 | 0.2291 | 0.1446 |
| $R_{free}$ | 0.1882 | 0.2642 | 0.2465 | 0.2605 | 0.1775 |
| *R.m.s.d values* | | | | | |
| Bond lengths (Å) | 0.006 | 0.014 | 0.001 | 0.003 | 0.007 |
| Bond angles (°) | 0.88 | 1.29 | 0.436 | 0.525 | 0.998 |
| *No. atoms* | | | | | |
| Protein | 728 | 728 | 1491 | 3126 | 724 |
| Ligand/ions | | 21 | 15 | 52 | 12 |
| *B-factors* (Å$^2$) | | | | | |
| Protein | 20.92 | 61.38 | 17.32 | 69.12 | 27.22 |
| ligand/ions | | 86.61 | 30.26 | 128.26 | 40.55 |
| *Ramachandran plot* | | | | | |

| | | | | | |
|---|---|---|---|---|---|
| Favored (%) | 100.0 | 95.6 | 98.8 | 97.7 | 100.0 |
| Allowed (%) | 0 | 4.4 | 1.2 | 2.3 | 0 |
| MolProbity score | 1.13 | 1.74 | 1.31 | 1.51 | 1.13 |
| Rotamer outliers (%) | 0.0 | 0.0 | 0.0 | 0.0 | 0.0 |
| *PDB code* | 6O8L | 6O8K | 6O8N | 6O8O | 6O8M |

*Highest-resolution shell values are shown in parentheses.

**Supplemental table 2.** Thermodynamic parameters for DNA binding with two cognate operators in SqrR variants obtained from fitting to a single-binding site model assuming tight dimer.

| | | | 200 mM NaCl[a] | 300 mM NaCl [a] | 400 mM NaCl [a] |
|---|---|---|---|---|---|
| Variant | Operator from promoter | Oxidation state | $K_{DNA}$ (x$10^9$ $M^{-1}$) | $K_{DNA}$ (x$10^9$ $M^{-1}$) | $K_{DNA}$ (x$10^9$ $M^{-1}$) |
| WT | 1451 | reduced | N.D. | 0.7 ± 0.1 | N.D. |
| C9S | 1451 | reduced | 21 ± 6 | 1.0 ± 0.2 | 0.19 ± 0.02 |
| | | No reducing agent | N.D. | 0.66 ± 0.05 | N.D. |
| | | disulfide | 0.02 ± 0.01 | N.D. | N.D. |
| | | tetrasulfide | 0.0039 ± 0.0002 | 0.0010± 0.0002 | N.D. |
| | 0785 | reduced | 4 ± 1 | N.D. | N.D. |

[a] 10 mM HEPES, pH 7.0, 2 mM EDTA.

**Supplemental table 3.** Kinetic parameters for reactions with persulfide donors for SqrR and BigR *

| Protein | Donor/ Conc’n | Mechanism | $P \rightarrow P^{SSH}$ | $P_X \rightarrow P^{SSSS}$ | $P_X \rightarrow P^{SSS}$ | $P_X \rightarrow P^{SS}$ | $P_X \rightarrow P^{SSSSS}$ |
|---|---|---|---|---|---|---|---|
| | | | $k_{PER}$ ($x10^{-3}$ s) | $k_{TET}$ ($x10^{-3}$ s) | $k_{TRI}$ ($x10^{-3}$ s) | $k_{DI}$ ($x10^{-3}$ s) | $k_{PENTA}$ ($x10^{-3}$ s) |
| C9S SqrR 0.03mM | GSSH 0.6 mM | 1 ($P_X$ =P) | - | 2.02 ± 0.07[a] | 0.18 ± 0.03 | 0.08 ± 0.03 | - |
| | GSSH 1.2 mM | 1 | - | 5.1 ± 0.4 | 0.034 ± 0.001 | 0.001 ± 0.001 | - |
| | | 2 ($P_X$ =$P^{SSH}$) | 6.8 ± 0.6 | 19 ± 4 | 1.1 ±0.6 | 0.4 ± 0.3 | - |
| | GSSH 0.6 mM + hemin 0.03 mM | 1 | - | 4.5 ± 0.5[a] | - | - | - |
| | | 2 | 6.6 ± 0.6 | 11 ± 2 | - | - | - |
| | GSSH 0.6 mM + $Cyt^{3+}$ 0.03 mM | 1 | - | 1.4 ± 0.8 | 0.06 ± 0.02 | 0.05 ± 0.02 | - |
| | GSSH 0.6 mM + DNA 0.03 mM | 2 | - | 0.27 ± 0.02 | 0.05 ± 0.01 | 0.03 ± 0.01 | - |
| | SNAP 0.6 mM | 1 | - | 0.58 ± 0.02 | 0.0084 ± 0.0001 | 0.004 ± 0.001 | - |
| BigR 0.03mM | GSSH 0.6 mM | 1 | - | 0.11 ± 0.01 | 0.003 ± 0.001 | 0.002 ± 0.001 | 0.36 ± 0.01 |
| | | 2 | 0.68 ± 0.04 | 0.7 ± 0.1 | 0.15 ± 0.06 | 0.009 ± 0.005 | 1.97 ± 0.04 |

[*] The data presented here are the result of the fitting presented in Figure 5B-C, Figure 6 (main text) and Figure S10. [a] These values were used to estimate the order for GSSH for the tetrasulfide formation to be 1.4 ± 0.2.

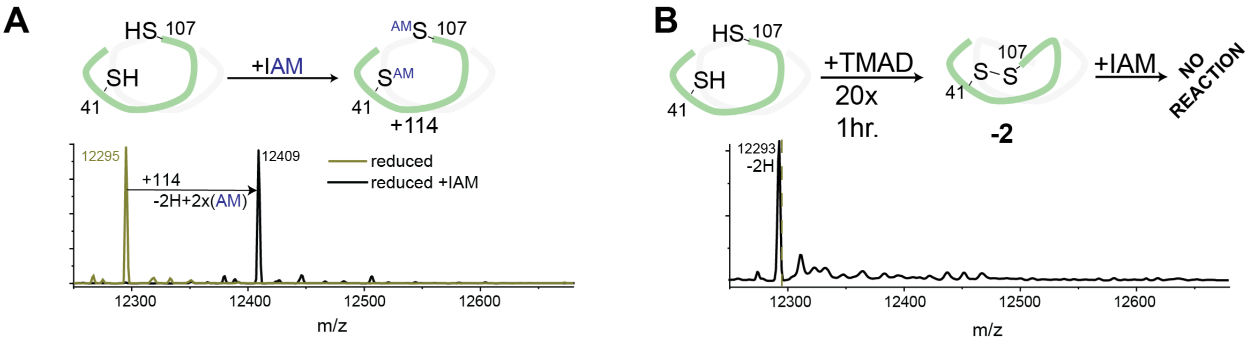


**Figure S1.** Illustration of quantitative capping of reduced C9S SqrR with IAM under native conditions and the reaction of C9S SqrR with TMAD. LC-ESI-MS spectra of (A) reduced and untreated C9S SqrR (*yellow* trace) and IAM treated C9S SqrR (*black* trace). (B) TMAD-treated C9S SqrR followed addition of an excess of IAM (900x). Solution conditions: 150 mM sodium phosphate, pH 7.4, 1 mM EDTA.

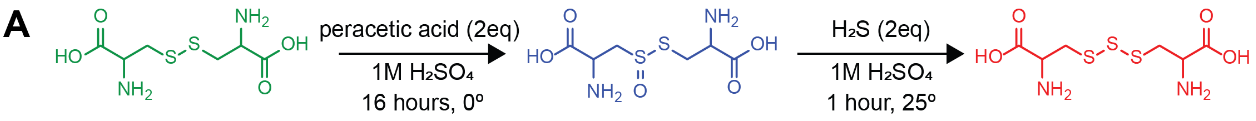


| Species | Cystine | Cystine-S-monoxide | Cysteine trisulfide |
|---|---|---|---|
| $[M+H]^+$ | 241.02 | 257.02 | 273.00 |
| $[M+Na]^+$ | 263.01 | 279.0 | 294.98 |
| $[M+2Na-H]^+$ | 284.90 | Not observed | Not observed |

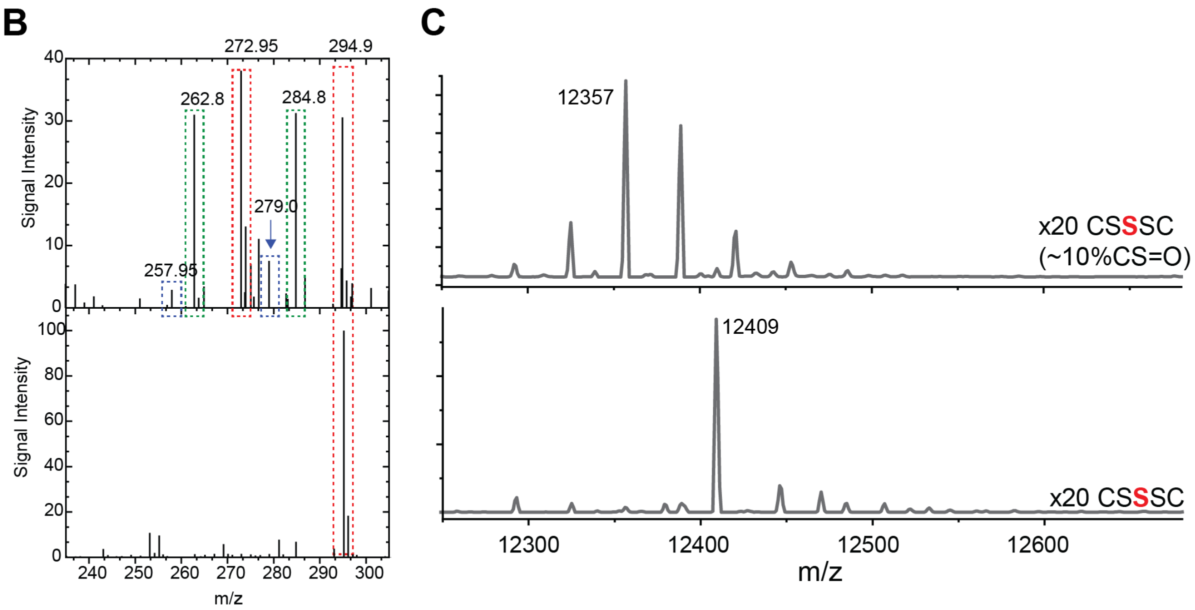


**Figure S2.** Cysteine trisulfide (CSSSC) synthesis and reaction with C9S SqrR. Reaction scheme for synthesis of CSSSC proceeding through a cystine-S-monoxide intermediate (*blue*) from cystine (*green*) starting material. Table includes the expected *m/z* for these species. (B) ESI-MS of impure (*top*) and pure (*bottom*) synthetic preparations of cysteine trisulfide. Peaks are highlighted for each molecule based on the color scheme in (A); masses given are the observed *m/z*. (C) LC-ESI-MS spectra of CSSSC-reacted C9S SqrR using 20-fold excess of impure (*top* trace) and pure (*bottom* trace) CSSSC, followed by addition of excess IAM. Solution conditiions: 30 µM protomer C9S SqrR,150 mM phosphate buffer pH 7.4, 1 mM EDTA.

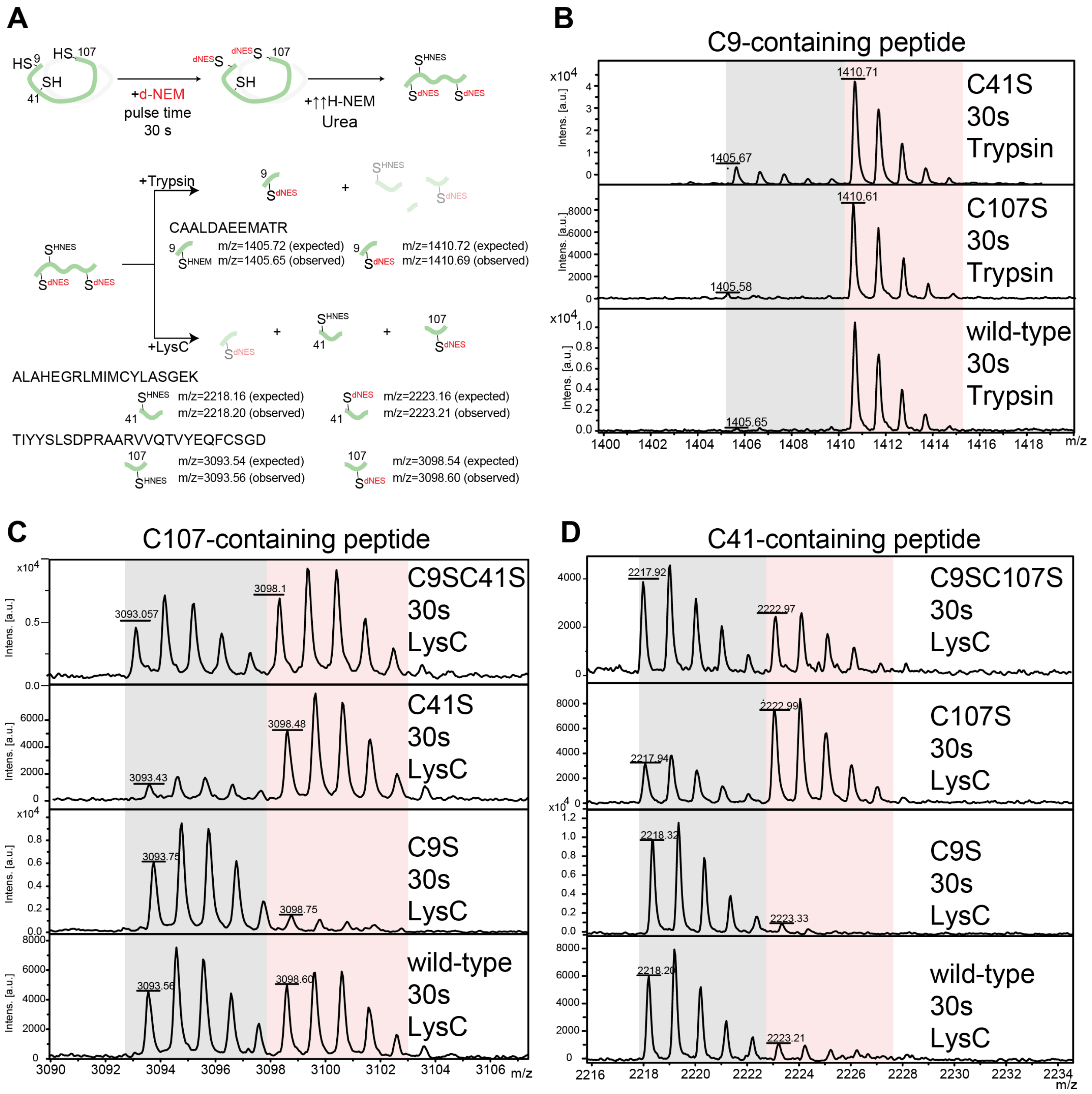


**Figure S3.** rPA-MS single time point assays used to compare the relative reactivies of C9, C41 and C107 in SqrR variants. (A) Reaction scheme exemplified for wild-type SqrR. Heavy *d5* (*grey* shaded) and light H5 (*red* shaded) C9- (B), C107- (C) and C41- (D) containing peptides for the different protein variants (indicated) at 30s pulse time with of 3-fold molar excess of reagent. C9 is more reactive than C107 (increased relative abundance of the "heavy" *d*5-NEM used for the alkylation pulse), which is significantly higher than C41 in all the variants tested. Substitution of the non-conserved C9 decreases the reactivity of the other Cys residues consistent with a slight protective role of the N-terminal unstructured (Figure S6) region in SqrR. Serine substitution of C41 or C107 increases the apparent nucleophilicity on the remaining Cys with little impact on C9. Solution conditions: 25 mM HEPES, pH 7.0, 200 mM NaCl, 5 mM EDTA.

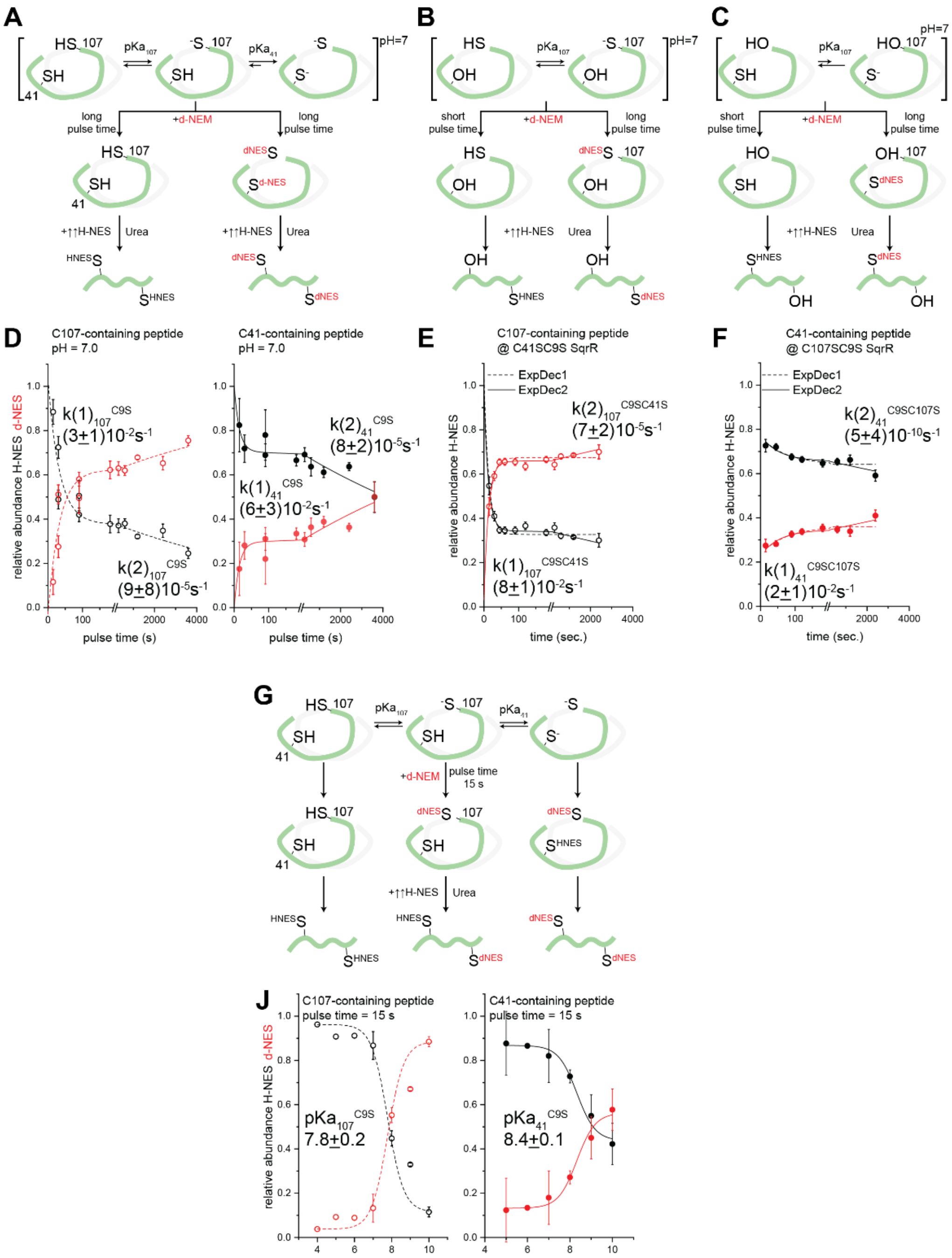


**Figure S4.** rPA-MS kinetics and estimation of the p$K_a$ of C41 and C107. Relative quantification of *d*5-NEM and H5-NEM alkylation events at C41 and C107 in the reduced state as a function of *d*5-NEM pulse time for C9S SqrR (A: reaction scheme; D: C107 and C41 containing peptides), C9S/C41S SqrR (B: reaction scheme; D: C107 containing peptide) and C9S/C107S (C: reaction scheme; D: C41 containing peptide). pH dependence of the relative quantification of *d*5-NEM and H5-NEM at a 15 s *d*5-NEM pulse time (G: reaction scheme; J: C107 and C41 containing peptides for C9S SqrR).

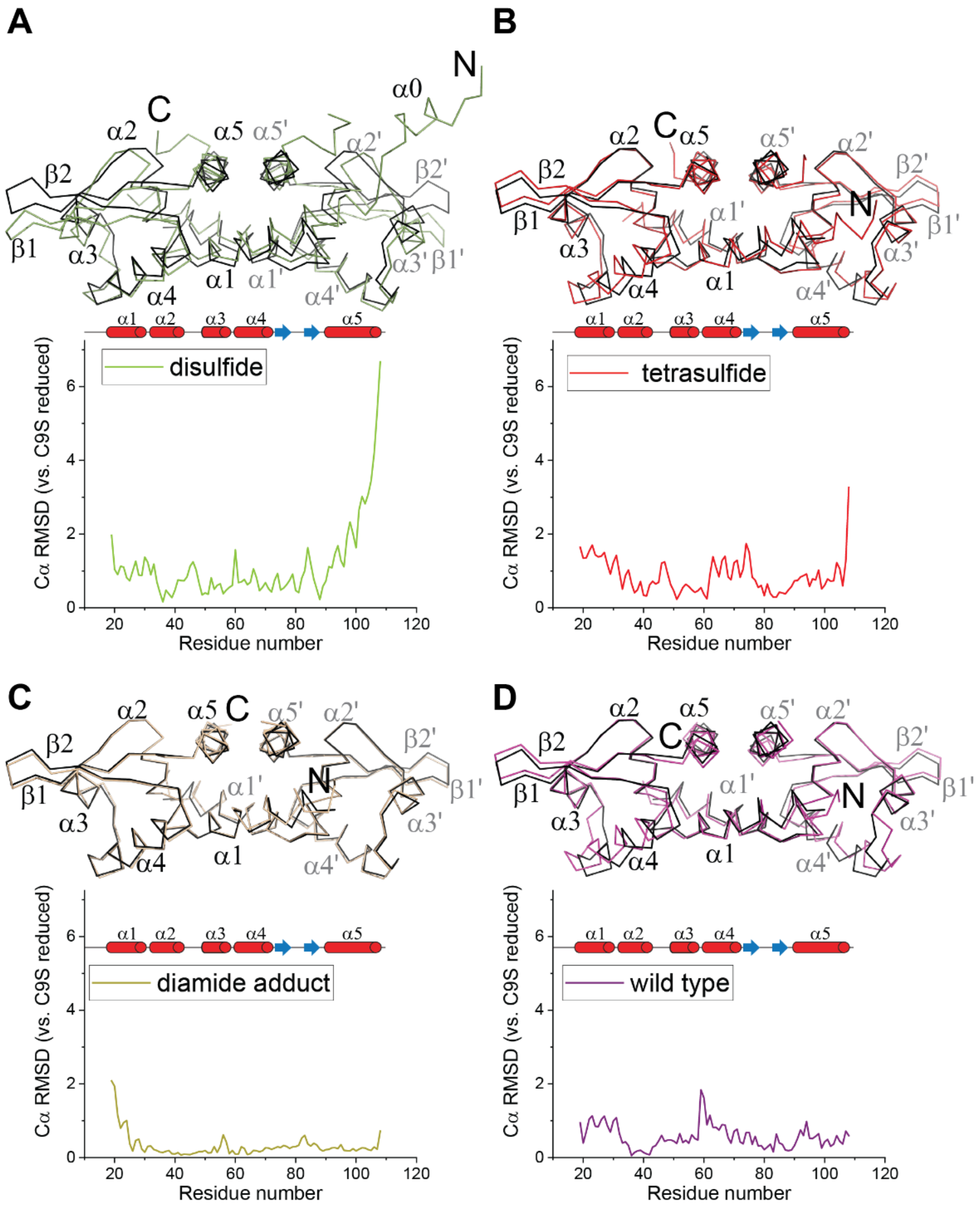


**Figure S5.** $C^{\alpha}$ ribbon diagram superpositions and pair-wise $C^{\alpha}$ root-mean square deviations (RMSD) in pairs of SqrR structures solved here. Superposition of the structures, depicted as a $C^{\alpha}$ trace, of reduced C9S SqrR with (A) disulfide crosslinked C9S SqrR; (B) tetrasulfide crosslinked C9S SqrR; (C) the C9S SqrR diamide adduct, and (D) wild-type reduced SqrR. The RMSD for $C^{\alpha}$ atoms for the monomer (calculated from the coordinates of chain A when there is more than one protomer in the unit cell) is shown at the bottom of each panel. The positions of the secondary structure elements are depicted as cylinder cartoons in each panel.

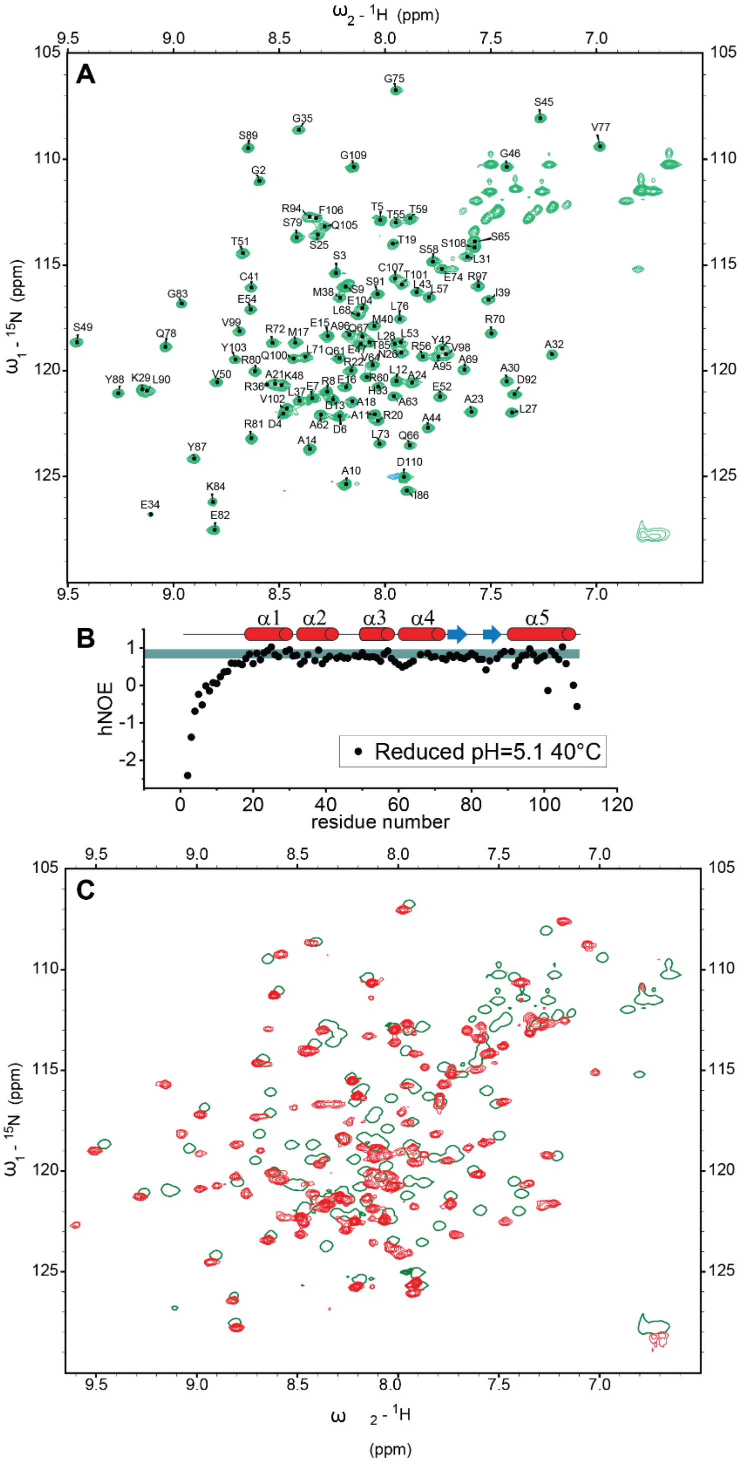


**Figure S6.** $^{1}$H, $^{15}$N HSQC spectra and $^{1}$H–{$^{15}$N} heteronuclear nuclear Overhauser enhancement (hNOE) of reduced C9S SqrR, (A) Resonance assigned $^{1}$H,$^{15}$N HSQC spectrum of C9S SqrR in the reduced state. Backbone assignments are shown in the one-letter code (assignments to be reported elsewhere). (B) Backbone hNOE as a function of residue number, indicating relative backbone flexibility of the N-terminal region (the shaded area indicates the expected values for a rigid backbone). The position of the secondary structure motifs is shown above. (C) $^{1}$H,$^{15}$N HSQC spectra of C9S SqrR in the tetrasulfide state compared to the reduced state (*green*

contour; single contour line shown only). The solution conditions for all the experiments were the same: 25 mM MES, 50 mM NaCl, pH 5.1, 40 °C).

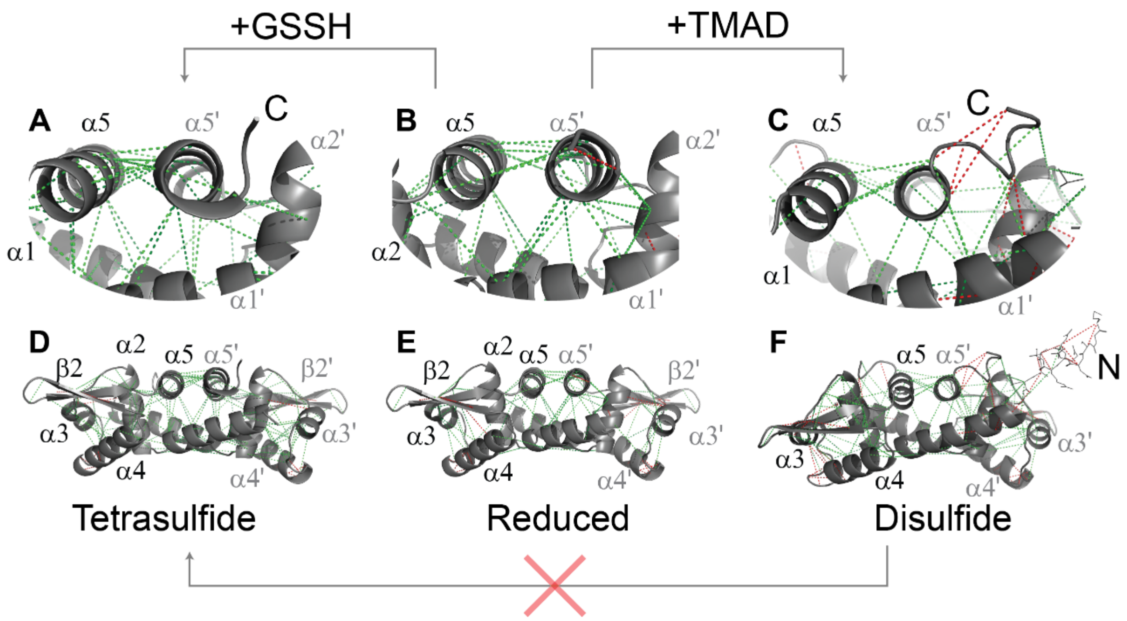


**Figure S7.** Frustration patterns in C9S SqrR in various allosteric states. Frustration patterns in C9S SqrR in the tetrasulfide (A, D), reduced (B, E) and disulfide states (C, F). The backbones of the proteins are shown as gray cartoons (the N-terminal region in the disulfide state is represented with lines), minimally frustrated contacts are depicted with *green* lines, and highly frustrated interactions with *red* lines. Neutral interactions were omitted for clarity. A higher density of highly frustrated contacts is observed in the vicinity of the disulfide bond (C), whereas another region of high local frustration for all three states of the protein is the N-terminal region of α4 (D-F).

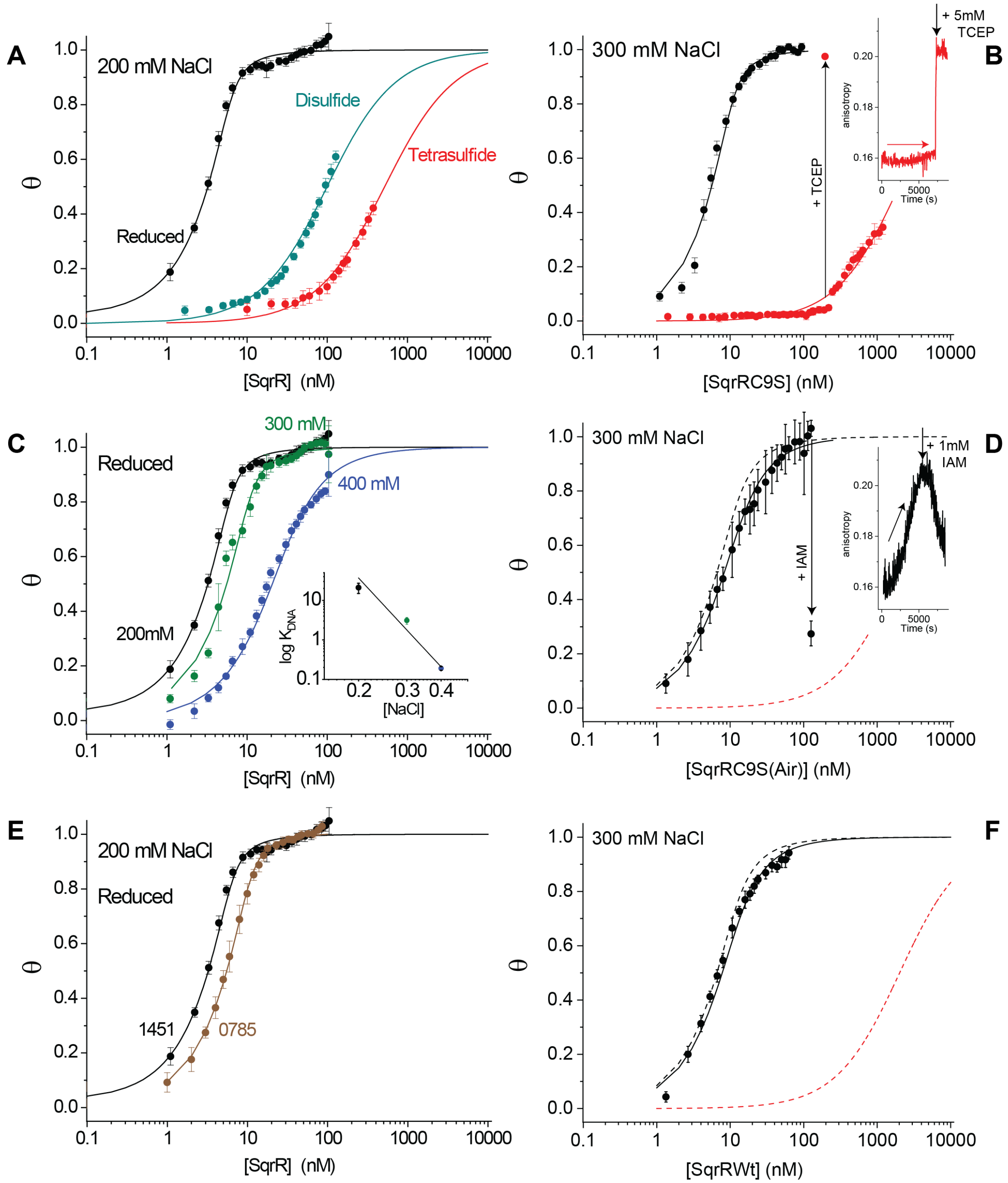


**Figure S8.** SqrO–DNA binding isotherms for C9S and wild-type SqrRs in the reduced, disulfide and tetrasulfide crosslinked states. (A) C9S SqrR in different oxidation states (200 mM NaCl), (B) C9S SqrR in the reduced and tetrasulfide states (300 mM NaCl), the arrow indicates addition of TCEP. The inset shows the anisotropy as a function of time for the titration of the protein followed by the addition of 5 mM TCEP that shows fast protein association after reduction. (C) C9S SqrR in the reduced state (2 mM TCEP added) at different [NaCl] (*inset*, [NaCl]-dependence of $K_a$). (D) C9S SqrR in the reduced state, no TCEP added (300 mM NaCl) (*black* continuous line) compared to C9S SqrR (reduced, *black* dashed lines; tetrasulfide, *red* dashed lines). The arrow indicates addition of IAM. The inset shows the anisotropy as a function of time for the titration of the protein followed by the addition of 1 mM IAM that shows a slow kinetic of dissociation. (E) C9S SqrR binding to the SqrO present in the promoter region of rcc01451 (*black*) and rcc0785 (brown) (300 mM NaCl), (F) Reduced wild-type SqrR (*black*) comparison with C9S SqrR (reduced, *black* dashed lines; tetrasulfide, *red* dashed lines) (300 mM NaCl). Conditions: 10 mM Hepes, pH 7.0, 25.0 °C, [SqrO]=5 nM rcc01451 unless indicated otherwise.

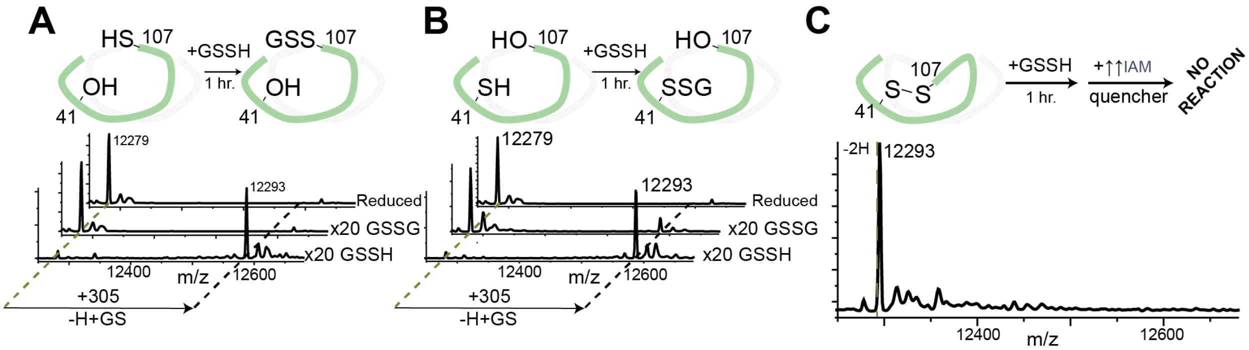


**Figure S9.** The reactivity of the putative persulfide intermediate in single Cys-containing SqrR variants and kinetic trapping of the disulfide. (A) LC-ESI-MS spectra of reduced C9S/C41S SqrR (30 μM protomer) treated for 1 h with a 20-fold molar excess of GSSG and GSSH. (B) LC-ESI-MS spectra of reduced C9S/C107S SqrR (30 μM protomer) treated for 1 h with a 20-fold molar excess of GSSG and GSSH. (C) LC-ESI-MS spectra of C9S SqrR in the disulfide state obtained following a 1 h incubation with 1 mM TMAD and reacted with a 20-fold molar excess of GSSH, followed by a large excess of IAM (900x) to cap. Solution conditions: 150 mM sodium phosphate, pH 7.4, 1 mM EDTA.

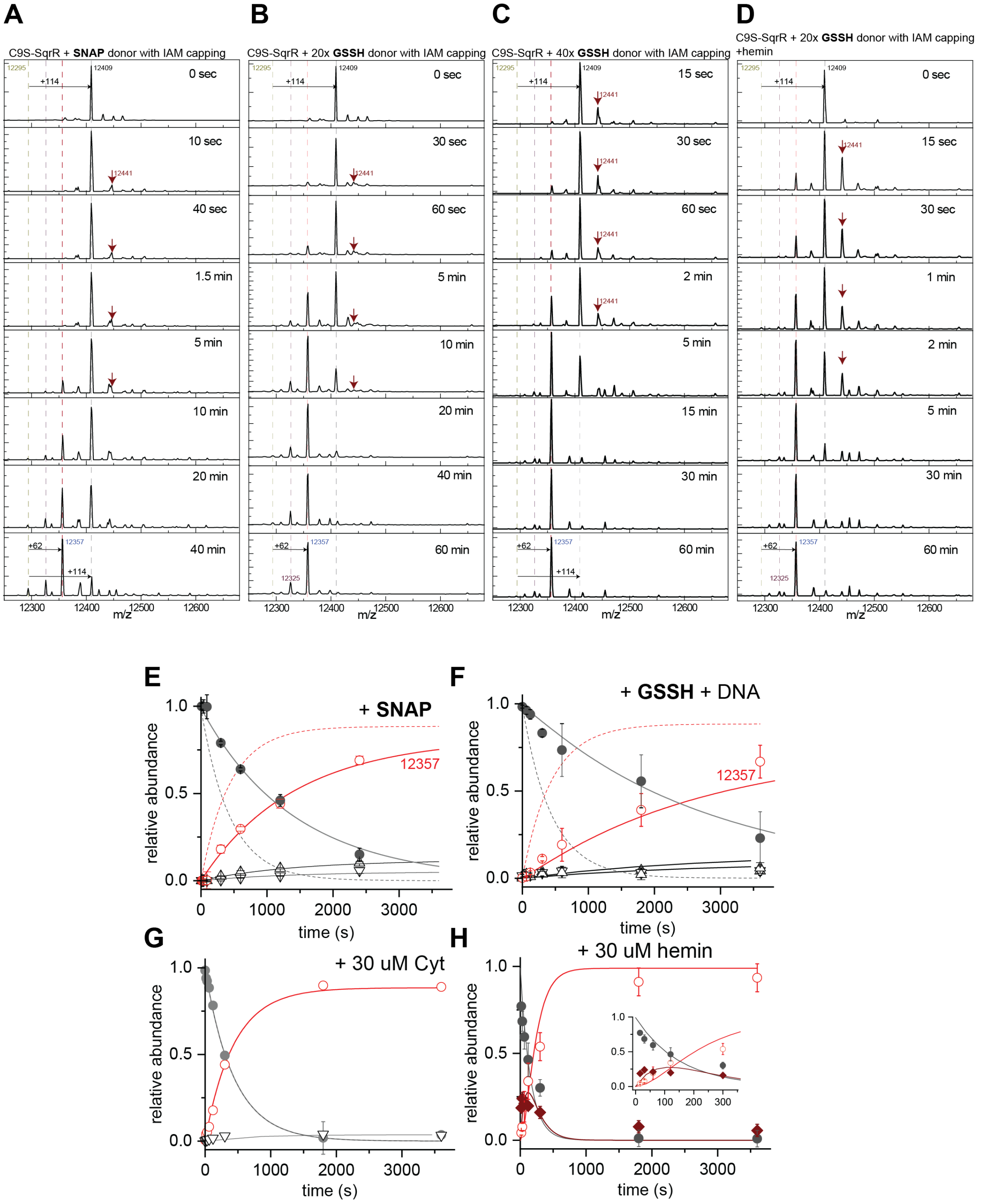


**Figure S10.** Representative kinetic traces that illustrate the time-course of reactivity of C9S SqrR toward various oxidants under various conditions. LC-ESI-MS spectra of C9S SqrR (30 μM) treated with a 20-fold excess SNAP donor (A), a 20-fold molar excess of GSSH (B), a 40-fold molar excess of GSSH (C) and a 20-fold molar excess GSSH with 30 μM hemin added (D) for variable times followed by a large excess of IAM (900x). Solution conditions: 150 mM sodium phosphate, pH 7.4, 1 mM EDTA. (E) Relative abundance of the different species as a function of pulse time obtained from the experiment in panel (A). (F) Relative abundance of the different species as a function of pulse time obtained from the experiment in panel (B), but carried out in the presence of 30 μM DNA operator (rcc1541). (G) Relative abundance of the different species as a function of pulse time obtained from the experiment in panel (B), but carried out in the presence of 30 μM oxidized cytochrome c (Cyt). (H) Relative abundance of the different species as a function of pulse time obtained from the experiment in panel (D) in the presence of 30 μM hemin. The continuous lines drawn through the data represent a nonlinear least squares fit to Mechanism (1) (panels E-G) or to Mechanism (2) (panel H).

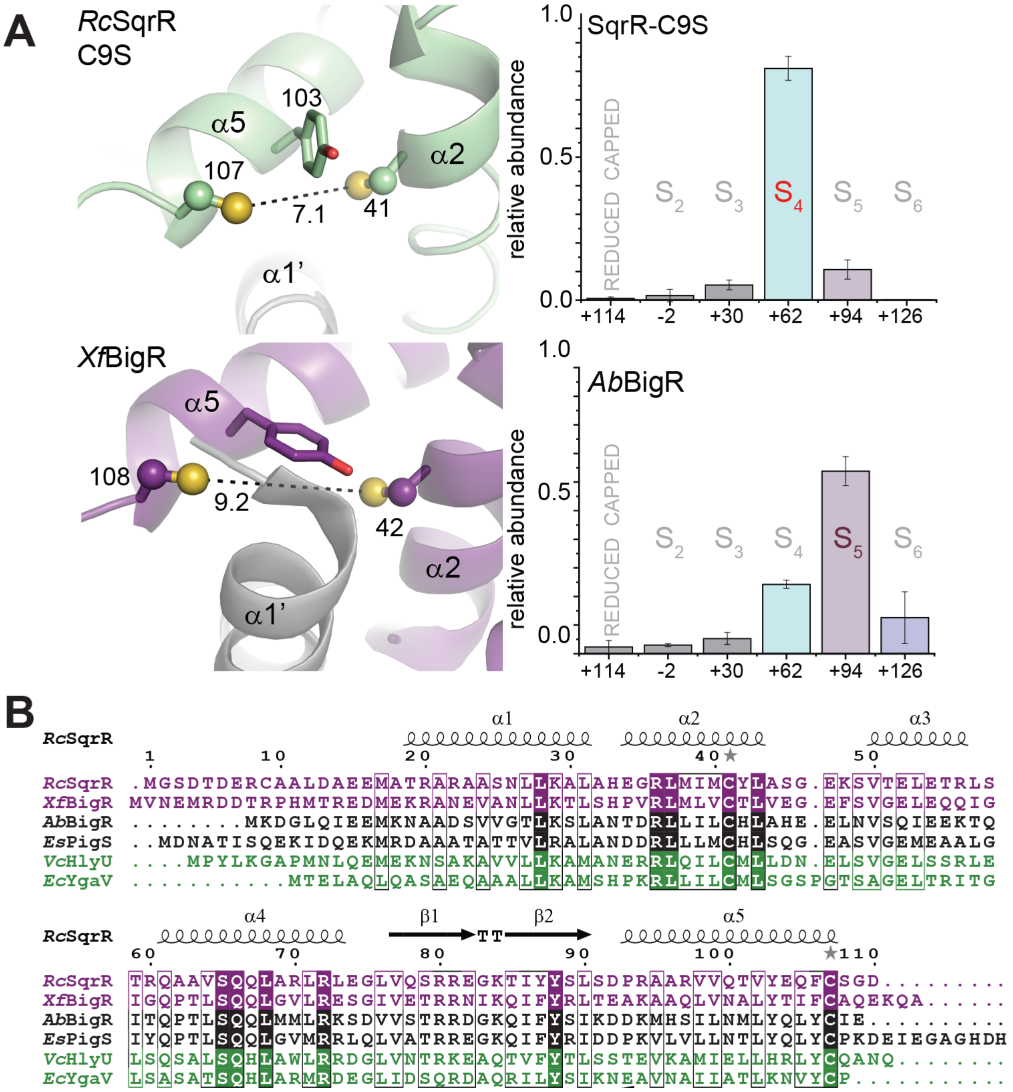


**Figure S11.** Comparison of the structures and polysulfide product distributions of *Rc* SqrR and BigR-like RSS sensors. (A) The cysteine pocket of C9S SqrR illustrating the distance between the thiols compared to that of *Xf* BigR in the reduced form (PDB 3pqj)[5]. The *Xf* BigR shows a folded N-terminal helix (labeled α1') that packs against the Cys cavity preventing the two Cys from getting closer to one another. (B) *Ab* BigR polysulfide length comparison with C9S SqrR showing that the pentasulfide is prevalent in *Ab* BigR while the tetrasulfide is the dominant product for SqrR. This suggests that the persulfide length can be determined by the initial distance between the two reduced thiolate Sγ atoms, supporting the idea that the final polysulfide distribution is dictated that which gives rise to a minimal structural perturbation. (B) Multiple sequence alignment of SqrR/BigR-like proteins that are characterized to distinct degrees (*purple*: *in vivo* evidence for persulfide specificity[1,23], *black*: persulfide reactivity can be inferred on the basis of the function of the regulated genes[24] (Walsh *et al*., manuscript submitted for publication); *green*: no information of the nature of the inducer is currently available[25,26]).

**REEFERENCES**


(1) Shimizu, T.; Shen, J.; Fang, M.; Zhang, Y.; Hori, K.; Trinidad, J. C.; Bauer, C. E.; Giedroc, D. P.; Masuda, S.; Bauer, C. E.; et al. *Proc. Natl. Acad. Sci.* **2017**, *114*, 2355.
(2) Busenlehner, L. S.; Cosper, N. J.; Scott, R. A.; Rosen, B. P.; Wong, M. D.; Giedroc, D. P. *Biochemistry* **2001**, *40*, 4426.
(3) Arunkumar, A. I.; Pennella, M. A.; Kong, X.; Giedroc, D. P. *Biomol. NMR Assign.* **2007**, *1*, 99.
(4) Liu, T.; Ramesh, A.; Ma, Z.; Ward, S. K.; Zhang, L.; George, G. N.; Talaat, A. M.; Sacchettini, J. C.; Giedroc, D. P. *Nat. Chem. Biol.* **2007**, *3*, 60.
(5) Guimarães, B. G.; Barbosa, R. L.; Soprano, A. S.; Campos, B. M.; De Souza, T. a.; Tonoli, C. C. C.; Leme, A. F. P.; Murakami, M. T.; Benedetti, C. E. *J. Biol. Chem.* **2011**, *286*, 26148.
(6) Adams, P. D.; Afonine, P. V.; Bunkóczi, G.; Chen, V. B.; Davis, I. W.; Echols, N.; Headd, J. J.; Hung, L. W.; Kapral, G. J.; Grosse-Kunstleve, R. W.; et al. *Acta Crystallogr. Sect. D Biol. Crystallogr.* **2010**, *66*, 213.
(7) Emsley, P.; Lohkamp, B.; Scott, W. G.; Cowtan, K. *Acta Crystallogr. D. Biol. Crystallogr.* **2010**, *66*, 486.
(8) Chen, V. B.; Arendall, W. B.; Headd, J. J.; Keedy, D. A.; Immormino, R. M.; Kapral, G. J.; Murray, L. W.; Richardson, J. S.; Richardson, D. C. *Acta Crystallogr. Sect. D Biol. Crystallogr.* **2010**, *66*, 12.
(9) Parra, R. G.; Schafer, N. P.; Radusky, L. G.; Tsai, M. Y.; Guzovsky, A. B.; Wolynes, P. G.; Ferreiro, D. U. *Nucleic Acids Res.* **2016**, *44*, W356.
(10) Freiberger, M. I.; Guzovsky, A. B.; Wolynes, P. G.; Parra, R. G.; Ferreiro, D. U. *Proc. Natl. Acad. Sci.* **2019**, *116*, 4037.
(11) Lindström, I.; Dogan, J. *ACS Chem. Biol.* **2018**, *13*, 1218.
(12) Ferreiro, D. U.; Hegler, J. A.; Komives, E. A.; Wolynes, P. G. *Proc. Natl. Acad. Sci. U. S. A.* **2011**, *108*, 3499.
(13) Kabil, O.; Banerjee, R. *J. Biol. Chem.* **2012**, *287*, 44561.
(14) Bianco, C. L.; Akaike, T.; Ida, T.; Nagy, P.; Bogdandi, V.; Toscano, J. P.; Kumagai, Y.; Henderson, C. F.; Goddu, R. N.; Lin, J.; et al. *Br. J. Pharmacol.* **2019**, *176*, 671.
(15) Savige, W. E.; Eager, J.; Maclaren, J. A.; Roxburgh, C. M. *Tetrahedron Lett.* **1964**, *5*, 3289.
(16) Apuy, J. L.; Busenlehner, L. S.; Russell, D. H.; Giedroc, D. P. *Biochemistry* **2004**, *43*, 3824.
(17) Glauninger, H.; Zhang, Y.; Higgins, K. A.; Jacobs, A. D.; Martin, J. E.; Fu, Y.; Coyne Rd,

H. J.; Bruce, K. E.; Maroney, M. J.; Clemmer, D. E.; et al. *Chem. Sci.* **2018**, *9*, 105.
(18) Kuzmic, P. *Anal. Biochem.* **1996**, *237*, 260.
(19) Yadav, P. K.; Xie, P.; Banerjee, R. *J. Biol. Chem.* **2012**, *287*, 37611.
(20) Vitvitsky, V.; Miljkovic, J. L.; Bostelaar, T.; Adhikari, B.; Yadav, P. K.; Steiger, A. K.; Torregrossa, R.; Pluth, M. D.; Whiteman, M.; Banerjee, R.; et al. *ACS Chem. Biol.* **2018**, *13*, 2300.
(21) Hilser, V. J.; Anderson, J. a; Motlagh, H. N. *Proc. Natl. Acad. Sci.* **2015**, *112*, 11430.
(22) Capdevila, D. A.; Braymer, J. J.; Edmonds, K. A.; Wu, H.; Giedroc, D. P. *Proc. Natl. Acad. Sci. U. S. A.* **2017**, *114*, 4424.
(23) de Lira, N. P. V.; Pauletti, B. A.; Marques, A. C.; Perez, C. A.; Caserta, R.; de Souza, A. A.; Vercesi, A. E.; Paes Leme, A. F.; Benedetti, C. E. *Sci. Rep.* **2018**, *8*, 3508.
(24) Gristwood, T.; McNeil, M. B.; Clulow, J. S.; Salmond, G. P. C.; Fineran, P. C. *J. Bacteriol.* **2011**, *193*, 1076.
(25) Mukherjee, D.; Datta, A. B.; Chakrabarti, P. *Biochim. Biophys. Acta* **2014**, *1844*, 2346.
(26) Gueuné, H.; Durand, M.-J.; Thouand, G.; DuBow, M. S. *Appl. Environ. Microbiol.* **2008**, *74*, 1954.
